# Release of VAMP5-positive extracellular vesicles by retinal Müller glia *in vivo*

**DOI:** 10.1101/2022.04.20.488918

**Authors:** Valerie Demais, Anne Pohl, Kirsten A. Wunderlich, Anna M. Pfaller, Lew Kaplan, Amelie Barthélémy, Robin Dittrich, Berta Puig, Bernd Giebel, Stefanie M. Hauck, Frank W. Pfrieger, Antje Grosche

## Abstract

Cell-cell interactions in the central nervous system are based on the release of molecules mediating signal exchange and providing structural and trophic support through vesicular exocytosis and the formation of extracellular vesicles. The specific mechanisms employed by each cell type in the brain are incompletely understood. Here, we explored the means of communication used by Müller cells, a type of radial glial cells in the retina, which forms part of the central nervous system. Using immunohistochemical, electron microscopic, and molecular analyses, we provide evidence for the release of distinct extracellular vesicles from endfeet and microvilli of retinal Müller cells in adult mice *in vivo*. We identify VAMP5 as a Müller cell-specific SNARE component that is part of extracellular vesicles and responsive to ischemia, and we reveal differences between the secretomes of immunoaffinity-purified Müller cells and neurons *in vitro*. Our findings suggest extracellular vesicle-based communication as an important mediator of cellular interactions in the retina.

## Introduction

Glial cells control the development (Araujo et al., 2019; Lago-Baldaia et al., 2020; Tan et al., 2021) and function (García-Cáceres et al., 2019; Kofuji and Araque, 2021; Nave and Werner, 2021) of neurons, and they influence the outcome of pathologic conditions due to disease or injury (Kim et al., 2020; Linnerbauer et al., 2020; Patel et al., 2019; Raiders et al., 2021; Wilton et al., 2019; Wilton and Stevens, 2020). During the last years, much has been learned about glia-neuron interactions. Neurons and different types of glial cells communicate via intercellular contacts and via the release of molecules fulfilling multiple functions that range from structural, energy and trophic support to cell signaling (Giaume et al., 2021; Illes et al., 2019; Jha and Morrison, 2020; Seifert and Steinhäuser, 2018; Shen et al., 2017; Sultan et al., 2015). Molecules of the secretome that are contained in intracellular vesicles can be released directly into the extracellular space following fusion of the vesicular membrane with the plasma membrane (Fiacco and McCarthy, 2018; Murat and García-Cáceres, 2021; Savtchouk and Volterra, 2018; Vardjan et al., 2019; Mielnicka and Michaluk, 2021). Alternatively, they can be released as molecular assemblies contained in extracellular vesicles (EVs) harboring a context- and cell type-specific host of proteins, lipids, and nucleic acids (Kalluri and LeBleu, 2020; Mathieu et al., 2019; Pfrieger and Vitale, 2018; Pistono et al., 2020; Théry et al., 2018; van Niel et al., 2022; Yates et al., 2022). EVs have been discussed as potential means of cell-cell communication in the normal and diseased central nervous system (CNS) (Aires et al., 2021; Budnik et al., 2016; Li et al., 2019; Liu et al., 2020; Lizarraga-Valderrama and Sheridan, 2021; Mahjoum et al., 2021; Pascual et al., 2020; Schnatz et al., 2021; You and Ikezu, 2019), but knowledge about their presence, origin, and composition in nervous tissues *in vivo* is still incomplete (Brenna et al., 2021). Here, we explored possible mechanisms of glial communication in the CNS using the mouse retina as model tissue. We focus on Müller cells, a prominent type of radial glial cell that spans the entire retina (Wang et al., 2017) and impacts retinal development and function by diverse mechanisms (Reichenbach and Bringmann, 2020). Previous studies suggested that these cells employ vesicular release of molecules (Slezak et al., 2012; Wagner et al., 2017), but the full range of their intercellular communication capacity is unknown. Our findings suggest that Müller cells release EVs from their endfeet facing the vitreous body and their microvilli surrounding photoreceptor segments. We show that Müller cell-derived EVs bear a characteristic protein composition that differs substantially from those secreted by neurons. Moreover, we uncover that in retinae of adult mice, vesicle-associated membrane protein 5 (VAMP5), a component of soluble N-ethylmaleimide-sensitive factor attachment proteins receptor (SNARE) complexes, is specifically expressed by Müller cells and contained in a subset of their EVs.

## Materials and Methods

### Animals

All experiments were performed with adult C57BL/6J mice in accordance with the European Community Council Directive 2010/63/EU and the ARVO Statement for the Use of Animals in Ophthalmic and Vision Research. Animals were housed in a 12h light/dark cycle with ∼400 lux with *ad libitum* access to drinking water and food.

### Transient retinal ischemia

The protocols for induction of transient retinal ischemia were approved by the local Bavarian authorities (55.2 DMS-2532-2-182, Germany). Ischemia was induced in one eye of 8-weeks-old male and female mice using the high intraocular pressure (HIOP) method (Pannicke et al., 2014; Wagner et al., 2016). The untreated contralateral eye served as internal control. This approach reduced the number of animals used in these experiments as demanded by the 3R rules. Anesthesia was induced by intraperitoneal injection of ketamine (100 mg/kg body weight; Ratiopharm, Ulm, Germany) and xylazine (5 mg/kg; Bayer Vital, Leverkusen, Germany). Pupillary dilation was induced by atropine sulfate (100 mg/kg; Braun, Melsungen, Germany). The anterior chamber of the test eye was cannulated from the pars plana with a 30-gauge infusion needle connected to a bottle containing saline (0.9% NaCl). The intraocular pressure was increased transiently to 160 mm Hg by elevating the bottle for 90 min. After removing the needle, animals were returned to cages and sacrificed after indicated periods of time for tissue analyses using carbon dioxide.

### Immunoaffinity-based cell purification

Specific cell types were enriched from retinal cell suspensions as described previously using immunomagnetic separation (Grosche et al., 2016; Pauly et al., 2019). Briefly, retinae were treated with papain (0.2 mg/ml; Roche Molecular Biochemicals) for 30 min at 37°C in the dark in calcium- and magnesium-free extracellular solution (140 mM NaCl, 3 mM KCl, 10 mM HEPES, 11 mM glucose, pH 7.4). After several rinses and 4 min of incubation with DNase I (200 U/ml), retinae were triturated in extracellular solution with 1 mM MgCl_2_ and 2 mM CaCl_2_ added. The retinal cell suspension was subsequently incubated with CD11b- and CD31-binding microbeads to remove microglia and vascular cells, respectively, according to the manufacturer’s instructions (Miltenyi Biotec, Bergisch Gladbach, Germany). The respective binding cells were depleted from the retinal suspension using large cell (LS)-columns, prior to Müller cell enrichment. To select Müller cells, the cell suspension was incubated in extracellular solution containing biotinylated anti-CD29 (0.1 mg/ml, Miltenyi Biotec) for 15 min at 4°C. Cells were washed in extracellular solution, spun down, resuspended in the presence of anti-biotin MicroBeads (1:5; Miltenyi Biotec) and incubated for 15 min at 4°C. After washing, CD29-positive Müller cells were enriched using LS columns according to the manufacturer’s instructions (Miltenyi Biotec). Cells in the flow-through of the last sorting step – depleted of microglia, vascular cells, and Müller cells – were considered as mixed neuronal population.

### Immunohistochemical and -cytochemical staining

For immunohistochemical staining, enucleated eyes were immersion-fixed (4% paraformaldehyde for 2h), washed with phosphate-buffered saline (PBS), cryoprotected in sucrose, embedded in Tissue-Tek® O.C.T. compound (Sakura Finetek, Staufen, Germany), and cut in 20 µm sections using a cryostat. Retinal sections were permeabilized (0.3% Triton X-100 plus 1% DMSO in PBS) and blocked (5% normal goat serum with 0.3% Triton X-100 and 1% DMSO in PBS) for 2h at room temperature (RT). For immunocytochemical staining, acutely isolated or cultured cells were fixed (4% paraformaldehyde for 15 min), washed with PBS, permeabilized (0.3% Triton X-100 plus 1% DMSO in PBS) and blocked (5% normal goat serum with 0.3% Triton X-100 and 1% DMSO in PBS; 30 min at RT). Sections or fixed cells were incubated with primary antibodies (Supplementary Table 1) in bovine serum albumine (BSA; 1% in PBS) overnight at 4°C. Samples were washed (1% BSA) and incubated with secondary antibodies (2h at RT; 1% BSA). In some experiments, cell nuclei were labeled with DAPI (1:1000; Life Technologies). Samples were mounted in Aqua-Poly/Mount (Polysciences, Hirschberg, Germany). Control experiments without primary antibodies indicated absence of unspecific labeling except for the goat-anti-mouse secondary antibody that labeled blood vessels (not shown). Images of stained sections and cells were acquired with a custom-made VisiScope CSU-X1 confocal system (Visitron Systems, Puchheim, Germany) equipped with a high-resolution sCMOS camera (PCO AG, Kehlheim, Germany). For super resolution microscopy, fixed retinal sections were permeabilized with Triton X-100 (0.5 % in 2% BSA in PBS) for 2h before incubation with primary antibodies in blocking solution (overnight at 4°C). After washing with PBS, sections were incubated with secondary antibodies (Abberior Star) and DAPI for 2h in blocking solution at RT, subsequently washed with PBS, and briefly rinsed with distilled water before being mounted with ProLong Gold antifade reagent (Invitrogen, Life Technologies). Stimulated emission depletion (STED) microscopy was performed at the Core Facility Bioimaging of the Biomedical Center of the LMU München with an inverted Leica SP8 STED X WLL microscope using appropriate lasers for fluorophore excitation (405 nm; pulsed white light laser 470 – 670 nm). Images were acquired with a 93x/1.3 NA glycerol immersion objective with the pixel size set to 23 nm. The following spectral settings were used: DAPI (excitation: 405 nm; emission: 415 – 450 nm; PMT), AbberiorStar 580 (580 nm; 590 – 620 nm; HyD; depletion laser: pulsed 775 nm, at 12% intensity), AbberiorStar 635P (635 nm; 645 – 702 nm; HyD; depletion laser: pulsed 775 nm, at 12% intensity). Recording of color channels was performed sequentially to avoid bleed-through. Image brightness and contrast were adjusted with the open source software FIJI (Schindelin et al., 2012).

### Immunoblotting of proteins from acutely isolated, immunoaffinity-purified cells

Pellets of enriched cell populations from pooled pairs of mouse eyes were redissolved in reducing radioimmunoprecipitation assay (RIPA) buffer, denatured, and sonicated. Protein amounts were quantified using the Bradford or the RC DC Protein assays (BioRad, Feldkirchen, Germany) according to the manufacturer’s instructions. Equal amounts of protein per sample were loaded to compare levels of selected VAMPs in cells (2.5 or 5 µg depending on the VAMP). Samples were separated by 12% SDS-PAGE and immunodetection of proteins was performed as described (Schäfer et al., 2017) using primary and secondary antibodies (Supplementary Table 1) diluted in blocking solution (5% BSA in Tris-buffered saline with Tween 20). Blots were developed with WesternSure PREMIUM Chemiluminescent Substrate (LI-COR, Bad Homburg, Germany).

### Test of VAMP5 antibody specificity by immunoprecipitation

Retinae were dissected from adult mice and lysed mechanically and chemically in RIPA buffer. The protein concentration was determined by the RC DC Protein Assay (BioRad). For immunoprecipitation, to each sample (1 mg of total extract from 4 mouse retinae), 5 µg of anti-VAMP5 antibody or a rabbit IgG-control antibody was added (Supplementary Table 1) and the antibody-lysate mix was incubated on the rotator at 4°C for 4h. For affinity purification, 25 µl of Protein G Sepharose® (Merck, Darmstadt, Germany) were resuspended three times in TBS (0.5% NP40) and then centrifuged at 2,000g for 5 min. Beads were added to the antibody-lysate mix and incubated on the rotator at 4°C overnight. The complexes formed by antibody, antigen, and Protein G Sepharose® were pelleted (4,000g for 5 min) two times, and washed in TBS. Proteins were eluted in Laemmli buffer for 10 min at 37°C and spun down at 4,000g. Supernatant was collected and stored at −20°C until mass spectrometric analysis.

### Transcript profiling of acutely isolated, immunoaffinity-purified cells by RNAseq

Total RNA was isolated from pellets of enriched cell populations using the PureLink® RNA Micro Scale Kit (Thermo Fisher Scientific, Schwerte, Germany). Validation of RNA integrity and quantification of RNA concentrations were performed using the Agilent RNA 6000 Pico chip analyser according to the manufacturer’s instructions (Agilent Technologies, Waldbronn, Germany). Enrichment of mRNA, library preparation (Nextera XT, Clontech), quantification (KAPA Library Quantification Kit Illumina, Kapa Biosystems, Inc., Woburn, MA, USA) and sequencing on an Illumina platform (NextSeq 500 High Output Kit v2; 150 cycles) were performed at the service facility of the KFB Center of Excellence for Fluorescent Bioanalytics (Regensburg, Germany; www.kfb-regensburg.de). After de-multiplexing, at least 20 million reads per sample were detected. Quality control (QC) of the reads and quantification of transcript abundance were performed with the Tuxedo suite software (Langmead et al., 2009; Trapnell et al., 2009). To this end, the cutadapt routine was used to remove adapter sequences (Martin, 2011) and several QC measures were obtained with the fastqc routine. Next, the trimmed reads were aligned to the reference genome/transcriptome (GRCm38) with HISAT2 (Kim et al., 2015). Transcript abundance was estimated with the stringtie routine (Pertea et al., 2015) and expressed as fragments per kilobase pairs of transcripts per million reads (fragments per kilobase million, fpkm).

### Quantitative reverse-transcriptase polymerase chain reaction (qRT-PCR) of acutely isolated, immunoaffinity-purified cells

Total RNA was isolated from pellets of enriched cell populations using the PureLink® RNA Micro Scale Kit (Thermo Fisher Scientific, Schwerte, Germany). An on-column DNase digestion step (PureLink® DNase mixture, Thermo Fisher Scientific, for 20 min at RT) was included to remove genomic DNA (Roche Molecular Systems, Mannheim, Germany). RNA integrity was validated using the Agilent RNA 6000 Pico Assay according to the manufacturer’s instructions (Agilent Technologies, Waldbronn, Germany). First-strand cDNAs from the total RNA purified from each cell population were synthesized using the RevertAid H Minus First-Strand cDNA Synthesis Kit (Fermentas by Thermo Fisher Scientific, Schwerte, Germany). Primers were designed using the Universal ProbeLibrary Assay Design Center (Roche, Supplementary Table 2) and transcript levels of candidate genes were measured by qRT-PCR using the TaqMan hPSC Scorecard™ Panel (384 well, ViiA7, Life Technologies, Darmstadt, Germany) according to the manufacturer’s instructions.

### TUNEL staining of cultured cells

Immunoaffinity-purified cells were plated on coverslips, placed in 24-well plates and cultured under chemically defined, serum-free conditions in DMEM-F12 GlutaMax supplemented with Gibco™ Antibiotic-Antimycotic (1:100; Thermo Fischer Scientific) and NeuroBrews (1:50; Miltenyi Biotec). After 24h or 48h, cells were fixed (4% PFA, 15 min at RT), permeabilized (0.25% Triton® X-100 in PBS, 20 min at RT), washed twice with de-ionized water and pretreated for 10 min at RT in TdT reaction buffer (Click-iT™ Plus TUNEL Assay, Alexa Fluor™ 594 dye, ThermoFisher Scientific, Schwerte, Germany), before the enzyme mix was added and incubated for 60 min at 37°C. After two washes in 3% BSA/PBS (2 min each), the Click iT® reaction cocktail was added and incubated for another 30 min at RT. Subsequent staining for the Müller cell marker glutamine synthetase (GLUL) and DAPI co-staining was performed as described in the section ‘Immunohistochemical and - cytochemical staining’, after coverslips were washed two times (3% BSA/PBS for 5 min at RT). Images of cells were taken with a custom-made VisiScope CSU-X1 confocal system (Visitron Systems, Puchheim, Germany).

### Preparation of immunoaffinity-purified EVs from cultured cells

Approximately 600,000 and 300,000 immunomagnetically sorted neurons and Müller cells, respectively, were seeded on individual wells of a 48-well-plate and cultured under chemically defined, serum-free conditions in 400 µl DMEM-F12 GlutaMax supplemented with Gibco™ Antibiotic-Antimycotic (1:100; Thermo Fischer Scientific) and NeuroBrews (1:50; Miltenyi Biotec). Small samples of cells were fixed and air-dried on glass slides to control for cell purity by immunocytochemical staining of GLUL. After 48h to 60h, conditioned medium (CM) was collected from individual wells of culture plates and preserved for subsequent analyses of released material. As reference, cells were also harvested from individual wells, pelleted by centrifugation, and immediately lysed in hot SDS (0.1 % in distilled water) for mass spectrometry or in RIPA buffer for immunoblotting and frozen at −80°C until further processing. CM was cleared by several centrifugation steps with increasing speed and duration (600g for 10 min; 2,000g for 30 min; 10,000g for 45 min) as described (Théry et al., 2006). The supernatant of the last centrifugation step was subjected to immunomagnetic separation of secreted material positive for the surface markers CD63 and CD9 (Miltenyi) following the manufacturer’s instructions. Immunoaffinity-purified material containing EVs was characterized at the molecular and structural level according to the MISEV2018 guidelines (Théry et al., 2018). For protein analyses, proteins in material enriched from CM by immunomagnetic separation were directly eluted and lysed from the µ-columns still in the magnet either with hot SDS (0.1 % in distilled water) for mass spectrometry or in RIPA buffer for immunoblotting. The material in the flow-through of the washing steps was pelleted by ultracentrifugation (at 100,000g for 2h; Optima MAX-XP Ultracentrifuge from Beckman Coulter equipped with a TLA55 rotor) before lysis as described (Théry et al., 2006). Each lysed fraction was frozen at −80°C until further use.

### Immunoblotting of proteins from EVs, flow-through and corresponding cell lysates

Immunoaffinity-purified material from CM, flow-through and lysates of each cell type were diluted in Laemmli sample buffer, and separated using SDS-PAGE (15%). Semidry blotting was performed using TRISbase (2.5 mM with 192% glycine and 20% methanol) as transfer buffer. PVDF membranes were blocked using non-fat dry milk (5% in TBS with 0.1% Tween 20) and incubated with primary and secondary antibodies. Blots were developed with Clarity MaxTM Western ECL substrate (Bio-Rad Medical Diagnostics GmbH, Dreieich, Germany).

### Nanoparticle tracking analysis

EV-containing samples from CM were charactized by nanoparticle tracking analysis (NTA) allowing for particle quantification and average size estimation (Dragovic et al., 2011; Sokolova et al., 2011). The ZetaViewTM platform (ParticleMetrix, Meerbusch, Germany) was calibrated with polystyrene beads of 100 nm diameter (Thermo Fisher). Analyses were performed exactly as described (Görgens et al., 2019). Briefly, with 5 repetitions, videos were recorded at all 11 positions. The machine’s sensitivity was set to 75, the shutter to 75 and the framerate to 30.

### Quantitative proteomics - LC-MSMS analysis, label-free quantification and data analysis

Proteins from immunoaffinity-purified EVs, flow-through and lysates of cultured cells were proteolysed with LysC and trypsin with the filter-aided sample preparation procedure as described (Grosche et al., 2016; Wiśniewski et al., 2009). Acidified eluted peptides were analyzed on a Q Exactive HF-X mass spectrometer (Thermo Fisher Scientific) online coupled to an UItimate 3000 RSLC nano-HPLC (Dionex). Samples were automatically injected and loaded onto the C18 trap cartridge and after 5 min eluted and separated on the C18 analytical column (Acquity UPLC M-Class HSS T3 Column, 1.8 μm, 75 μm x 250 mm; Waters) by a 95 min non-linear acetonitrile gradient at a flow rate of 250 nl/min. MS spectra were recorded at a resolution of 60,000 with an automatic gain control (AGC) target of 3 x 10e6 and a maximum injection time of 30 ms from 300 to 1,500 m/z. From the MS scan, the 15 most abundant peptide ions were selected for fragmentation via HCD with a normalized collision energy of 28, an isolation window of 1.6 m/z, and a dynamic exclusion of 30s. MS/MS spectra were recorded at a resolution of 15,000 with a AGC target of 10e5 and a maximum injection time of 50 ms. Unassigned charges and charges of +1 and >+8 were excluded from precursor selection. Acquired raw data were analyzed in the Proteome Discoverer 2.4 SP1 software (Thermo Fisher Scientific; version 2.4.1.15) for peptide and protein identification via a database search (Sequest HT search engine) against the SwissProt Mouse database (Release 2020_02, 17061 sequences), considering full tryptic specificity, allowing for up to one missed tryptic cleavage site, precursor mass tolerance 10 ppm, fragment mass tolerance 0.02 Da. Carbamidomethylation of cysteine was set as a static modification. Dynamic modifications included deamidation of asparagine and glutamine, oxidation of methionine, and a combination of methionine loss with acetylation on protein N-terminus. The Percolator algorithm (Käll et al., 2007) was used to validate peptide spectrum matches and peptides.

Only top-scoring identifications for each spectrum were accepted, additionally satisfying a false discovery rate < 1% (high confidence). The final list of proteins satisfying the strict parsimony principle included only protein groups passing an additional protein confidence false discovery rate < 5% (target/decoy concatenated search validation). Quantification of proteins, after precursor recalibration, was based on intensity values (at RT apex) for all unique peptides per protein. Peptide abundance values were normalized to the total peptide amount. The protein abundances were calculated summing the abundance values for admissible peptides. To compare proteins in EVs and lysate, the final protein ratio was calculated using median abundance values of five biological replicates each. The statistical significance of the ratio change was tested with ANOVA. P values were adjusted for multiple testing by the Benjamini-Hochberg correction. Proteins with an adjusted P value < 0.05 were deemed significant. To generate the heatmaps using open source programming language R (Team, 2021), mass spectrometric data were first filtered to include only proteins that were detected in at least three of five biological replicates of the CD9- or C63-positive EV fractions of neurons or Müller cells. Moreover, proteins had to be enriched by 1.25 fold in the respective EV sample compared to the corresponding lysate. To generate an abundance heatmap, normalized abundance values were log2-transformed and the Manhattan distances between the samples and proteins, respectively, were calculated. Using hierarchical clustering with the ward.D2 method, dendrograms were generated that were separated in 11 clusters for the proteins and 2 clusters for the samples. The heatmap itself including annotations was created with the pheatmap package (https://cran.r-project.org/web/packages/pheatmap/index.html). The smaller 11 clusters were then combined into 4 major clusters, for which gene enrichment analysis was performed using g:Profiler (ordered query, g:SCS threshold of a p-value <0.05) (Raudvere et al., 2019). Venn diagrams were created using R (Team, 2021) with proteins showing a more than 2-fold increase in the respective CD9- or CD63-positive EV fraction compared to corresponding cell lysates prepared in parallel. Proteins were ranked based on abundance values in respective CD9- or CD63-positive EV fractions relative to values in cell lysates. To highlight established EV markers, protein data were downloaded from the Vesiclepedia website (version 4.1) (Pathan et al., 2019) and restricted to components that were detected by at least three independent experimental approaches.

### Immunogold labeling of retinae and transmission electron microscopy

Retinae were fixed (PFA 4%, glutaraldehyde 0.1% in PBS at pH 7.4 for 1h at RT), washed in PBS, incubated with saponin (0.1% in PBS for 1h), and washed with BSA (2% in PBS, five times for 10 min each). Retina were incubated overnight at 4°C with an antibody against VAMP5 (1:100), CD63 (1:50) or CD9 (1:50; 0.1% BSA in PBS). After washing (2% BSA in PBS), retinae were incubated with ultrasmall nanogold F(ab’) fragments of goat anti-rabbit or goat anti-mouse (1:100, 0.1% BSA in PBS; Aurion), rinsed (0.1% BSA in PBS) and then washed in PBS and water. Gold particles were silver-enhanced using the R-Gent SE-EM kit (Aurion, Wageningen, Netherlands). For double-immunogold labeling, retinae were washed again several times in distilled water and phosphate buffer (PB; 0.1M NaH_2_PO_4_; pH 7.4), blocked (5% normal goat serum in PBS) and incubated overnight at 4°C with an additional antibody. After washing (5% normal goat serum in PBS), retinae were incubated with ultrasmall nanogold F(ab’) fragments of goat anti-rat (1:100, 0.1% normal goat serum in PBS; Aurion). After several rinses in phosphate buffer, retina were fixed (2.5% glutaraldehyde in PB) and washed in PB and in distilled water. The second round of gold particles was silver enhanced using the R-Gent SE-EM kit (Aurion) until they showed distinct sizes from the first ones. Immunogold-labeled retinae were postfixed (0.5% OsO4 in distilled water) for 15 min. Finally, samples were dehydrated in graded ethanol series and embedded in Embed 812 (EMS). Ultrathin sections were cut with an ultramicrotome (Leica), stained with uranyl acetate [1% (w/v) in 50% ethanol], and examined by transmission electron microscopy (TEM; Hitachi H7500 equipped with an AMT Hamamatsu digital camera).

### Electron microscopic analysis of EVs secreted in vitro

To determine the size and shape of EV-like material secreted by cultured Müller cells and neurons, corresponding CM was subjected to differential ultracentrifugation (600g for 10 min; 2,000g for 30 min; 10,000g for 45 min; 100,000g for 2h) as described (Théry et al., 2006). Pellets of the last step were fixed (2% glutardaldehyde in PBS for 1h), dried overnight on glass coverslips (rinsed with distilled water, dehydrated in graded ethanol series, desiccated in hexamethyldisilazane, and air-dried before use), carbon-coated and examined by scanning electron microscopy (SEM) at 15kV (Hitachi S800). For electron microscopic inspection of immunogold-labeled EVs, immunoaffinity-purified material was fixed (4% PFA, 0.1% glutaraldehyde in PBS at pH 7.4 for 1h), washed (PBS), dried overnight on glass coverslips for scanning electron microscopy (SEM) or on formvar-carbon grids for TEM and washed (BSA 2% in PBS, five times for 10 min each). EV samples were incubated overnight at 4°C with antibodies against VAMP5 (1:100) and CD63 (1:50) or CD9 (1:50; 0.1% BSA in PBS), washed (2% BSA in PBS), and incubated for 1h with nanogold F(ab’) fragments (1:100 in PBS with 0.1% BSA, Aurion) with different particle sizes and secondary antibodies (goat anti-rabbit: 40 nm for SEM and 5 nm for TEM; goat anti-mouse or goat anti-rat: 25 nm for SEM or 15 nm for TEM). After several rinses in PB, EVs were fixed in glutaraldehyde (2.5% in PB), washed in PB and postfixed in OsO_4_ (0.5% in PB for 15 min). Finally, all samples were rinsed in distilled water, dehydrated in graded ethanol series, desiccated in hexamethyldisilazane, and air-dried. TEM samples were stained with uranyl acetate [1% (w/v) in distilled water] and examined (Hitachi H7500 equipped with an AMT Hamamatsu digital camera). For SEM analysis, immunogold-labeled samples were carbon-coated and examined at 15kV (Hitachi S800). Secondary electron and backscatter electron images were collected simultaneously by corresponding detectors.

### Statistical analysis

Statistical analyses were performed with indicated tests using GraphPad Prism software (version 7). In figures, significance levels are indicated by asterisks (*, p < 0.05; **, p < 0.01; ***, p < 0.001). Whiskers in figures indicate standard error of the mean.

## Results

We explored vesicle-based communication by retinal Müller cells using adult mice as experimental model. As a first step, we determined which vesicle-associated membrane proteins (VAMPs) are expressed by retinal Müller cells *in vivo* using immunohistochemical staining of retinal sections. We focused on the VAMP family as these proteins are part of the SNARE complex, which mediates membrane fusion and thereby enables the cellular release of molecules and vesicles (Südhof and Rothman, 2009; Urbina and Gupton, 2020).

### Expression of selected VAMPs in retinal Müller cells and their post-ischemic upregulation

As shown in Fig. 1a, each member of the VAMP family showed a distinct expression pattern in the adult mouse retina. Notably, we observed an overlap of cellubrevin/VAMP3, myobrevin/VAMP5 and VAMP8 with the Müller cell-specific markers GLUL or retinaldehyde binding protein 1 (RLBP1). On the other hand, VAMP1, 2, 7 were present in plexiform layers containing synaptic connections, whereas VAMP4 labeled cell somata in the ganglion cell and inner nuclear layer (Fig. 1a). We validated the glial expression of VAMP3, 5 and 8 using independent experimental approaches (Fig. 1b,c). Cell-specific transcript analyses by RNAseq and qRT-PCR using lysates of immunoaffinity-purified cells (Grosche et al., 2016; Pauly et al., 2019) corroborated our results obtained by immunohistochemical staining (Fig. 1b). Müller cells expressed *Vamp3*, *Vamp5* and *Vamp8*, whereas *Synaptobrevin/Vamp2*, a component of synapses, was enriched in neurons indicating the validity of cell-specific profiles. We also examined the presence of VAMP proteins in immunoaffinity-purified cells by immunoblotting (Fig. 1c) and its cellular distribution by immunocytochemical staining of acutely isolated Müller cells (Fig. 1d). These experiments revealed that Müller cells express VAMP3 and VAMP5 and that both VAMPs show a punctate distribution indicating their presence in vesicle-like structures (Fig. 1d). The validity of the VAMP5 antibody (Supplementary Table 1) was indicated by previously published observations that VAMP5-deficient mice show strongly reduced signals in immunoblots and after immunohistochemical staining performed with the same antibody (Ikezawa et al., 2018). Moreover, our work revealed VAMP5 as a top-enriched protein in immunoprecipitated retinal lysates analysed by mass spectrometry (Supporting Data 1).

**Figure 1.**
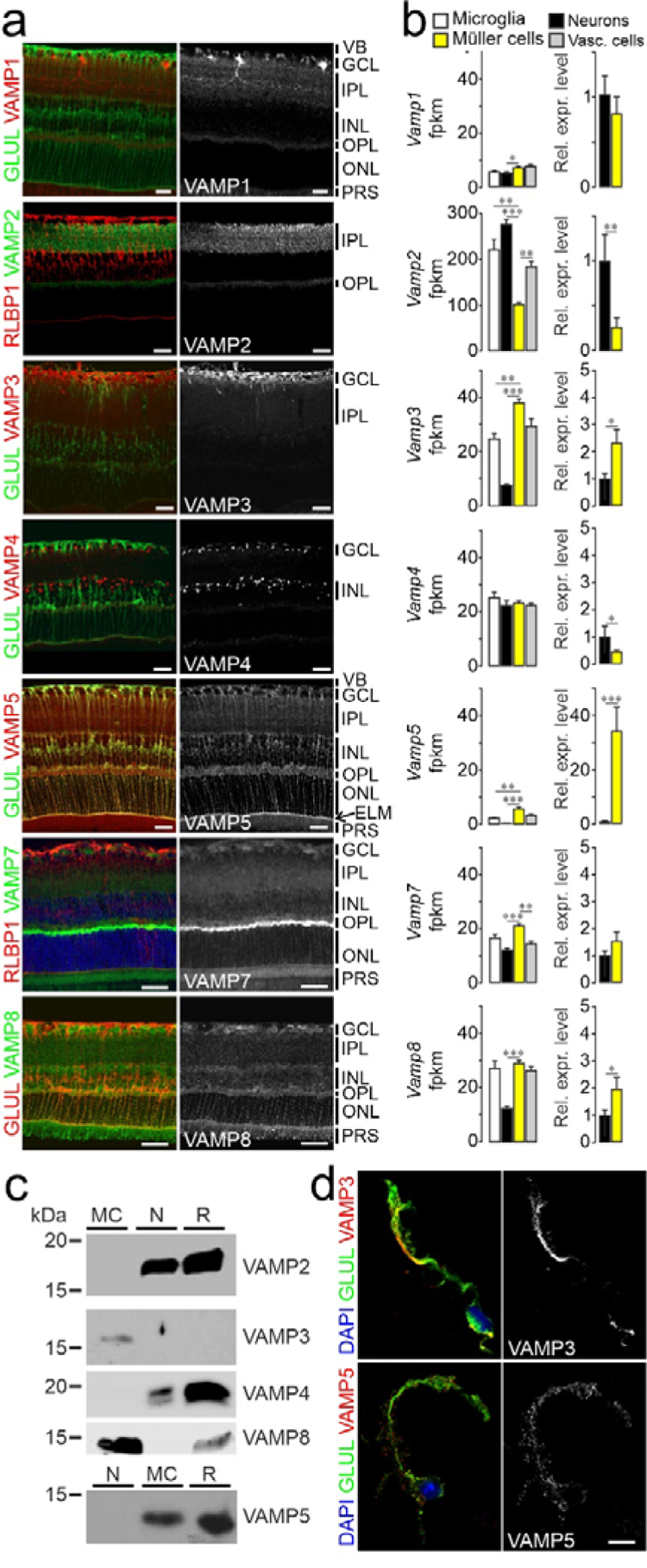
Expression of VAMP3, 5 and 8 by retinal Müller cells. (a) False-color and greyscale confocal micrographs of retinal sections from adult mice subjected to double-immunohistochemical staining of the indicated VAMPs and of the Müller cell markers GLUL and RLBP1. Positions of selected ocular structures and retinal layers are indicated: VB, vitreous body; GCL, ganglion cell layer; IPL, inner plexiform layer; INL, inner nuclear layer; OPL, outer plexiform layer; ONL, outer nuclear layer; ELM, external limiting membrane (arrow); PRS, photoreceptor segments. Scale bars: 20 µm. (b) Left, average counts of indicated *Vamp* transcripts obtained by RNAseq from lysates of indicated types of acutely isolated, immunoaffinity-purified retinal cells (n = 3-4 preparations; Mann-Whitney test; whiskers indicate SEM). Fpkm, fragments per kilobase million. Right, mean relative expression levels of indicated *Vamps* in Müller cells compared to neurons. (c) Representative immunoblots showing the presence of selected VAMPs in indicated populations of acutely isolated immunoaffinity-purified retinal cells (MC, Müller cells; N, neurons; R, retinal lysates). For each protein tested, equal amounts of total protein from indicated samples were loaded. (d) False-color and greyscale micrographs of acutely isolated immunoaffinity-purified Müller cells subjected to nuclear (blue) and double-immunocytochemical staining of VAMP3 and VAMP5 (red) and GLUL (green). Scale bar: 10 µm.

Glial cells react swiftly to injury and disease with prominent and context-specific changes in their expression profiles (Escartin et al., 2021; Pekny et al., 2016; Zamanian et al., 2012). To test whether expression levels of *Vamp3*, *5* and *8* in Müller cells change during gliosis, we used an established model of transient retinal ischemia (Pannicke et al., 2014; Wagner et al., 2016). We observed several-fold increases of these glia-specific *Vamp* transcripts in acutely isolated, immunoaffinity-purified Müller cells from ischemic retinae compared to cells from control tissue of contralateral eyes (Fig. 2a). Immunohistochemical staining revealed that transient ischemia enhanced levels of VAMP3 in processes of Müller cells in the plexiform layers and in structures located in the layer containing photoreceptor segments. VAMP5 levels rose in both plexiform layers and at the external limiting membrane. VAMP8 expression inceased across all plexiform layers. (Fig. 2b). Transcript levels of other members of the VAMP family were not altered. Taken together, these results showed that Müller cells express a subset of VAMPs, whose levels increase under pathologic conditions.

**Figure 2.**
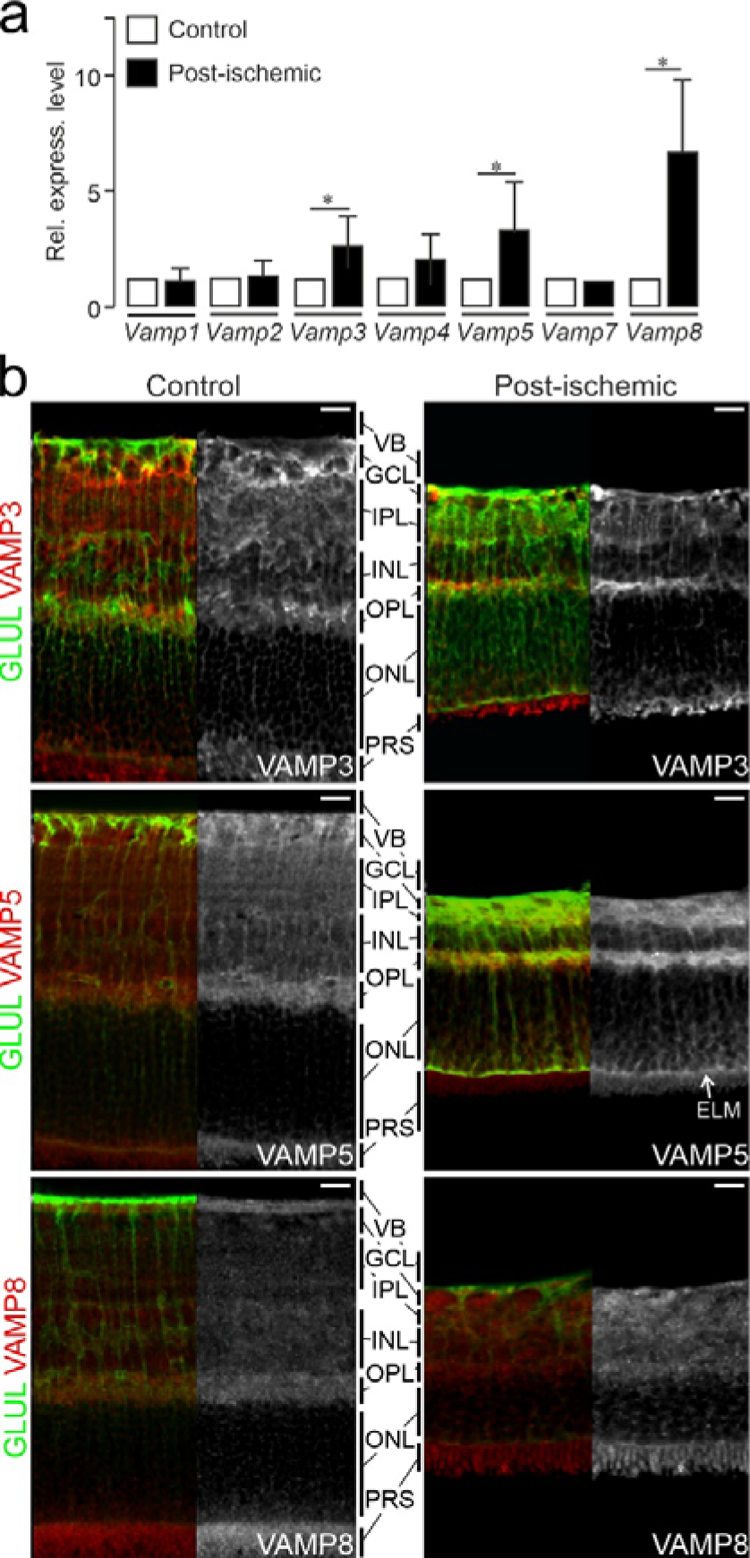
Ischemia-induced increase of *Vamp3*, *Vamp5* and *Vamp8* expression in retinal Müller cells. (a) Mean relative levels of indicated transcripts as determined by qRT-PCR in acutely isolated, immunoaffinity-purified Müller cells from retinae at 7 days after transient ischemia (post-ischemic). Levels were normalized to those in Müller cells isolated from untreated control eyes (n = 3 preparations; whiskers indicate SEM; Mann-Whitney test). (b) False-color and greyscale confocal micrographs of retinal sections from control eyes and from eyes at 7 days after transient ischemia followed by double-immunohistochemical staining of the indicated VAMPs (red) and of GLUL-positive Müller cells (green). VB, vitreous body; GCL, ganglion cell layer; IPL, inner plexiform layer; INL, inner nuclear layer; OPL, outer plexiform layer; ONL, outer nuclear layer; ELM, external limiting membrane; PRS, photoreceptor segments. Scale bar: 20 µm.

### Presence of VAMP5 in multivesicular bodies of retinal Müller cells and in the extracellular space facing the internal and external limiting membranes *in vivo*

The unexpectedly strong expression of VAMP5 in Müller cells prompted us to focus on this member of the VAMP family and to scrutinize its subcellular distribution using TEM. To facilitate comprehension of electron micrographs, morphologic features of Müller cells and their location with respect to the different retinal layers are shown schematically (Fig. 3a) together with representative transmission electron micrographs of their compartments in the inner and outer retina (Fig. 3b). Immunogold staining of retinal sections combined with TEM corroborated the presence of VAMP5 in Müller cells (Fig. 4). We detected the protein in distinct intracellular structures resembling vesicles and multivesicular bodies. These structures were located in endfeet of Müller cells and in their radial processes apposed to the external limiting membrane (ELM) of the outer retina (Fig. 4). Interestingly, we also observed VAMP5-positive vesicle-like structures on the extracellular side of the internal limiting membrane facing the vitreous body and around apical microvilli of Müller cells reaching into the subretinal space surrounding photoreceptor segments (Fig. 4). Together, our ultrastructural observations revealed the presence of VAMP5 in vesicular structures located in the cytoplasm of Müller cells and in the extracellular space facing the internal and external limiting membrane *in vivo*.

**Figure 3.**
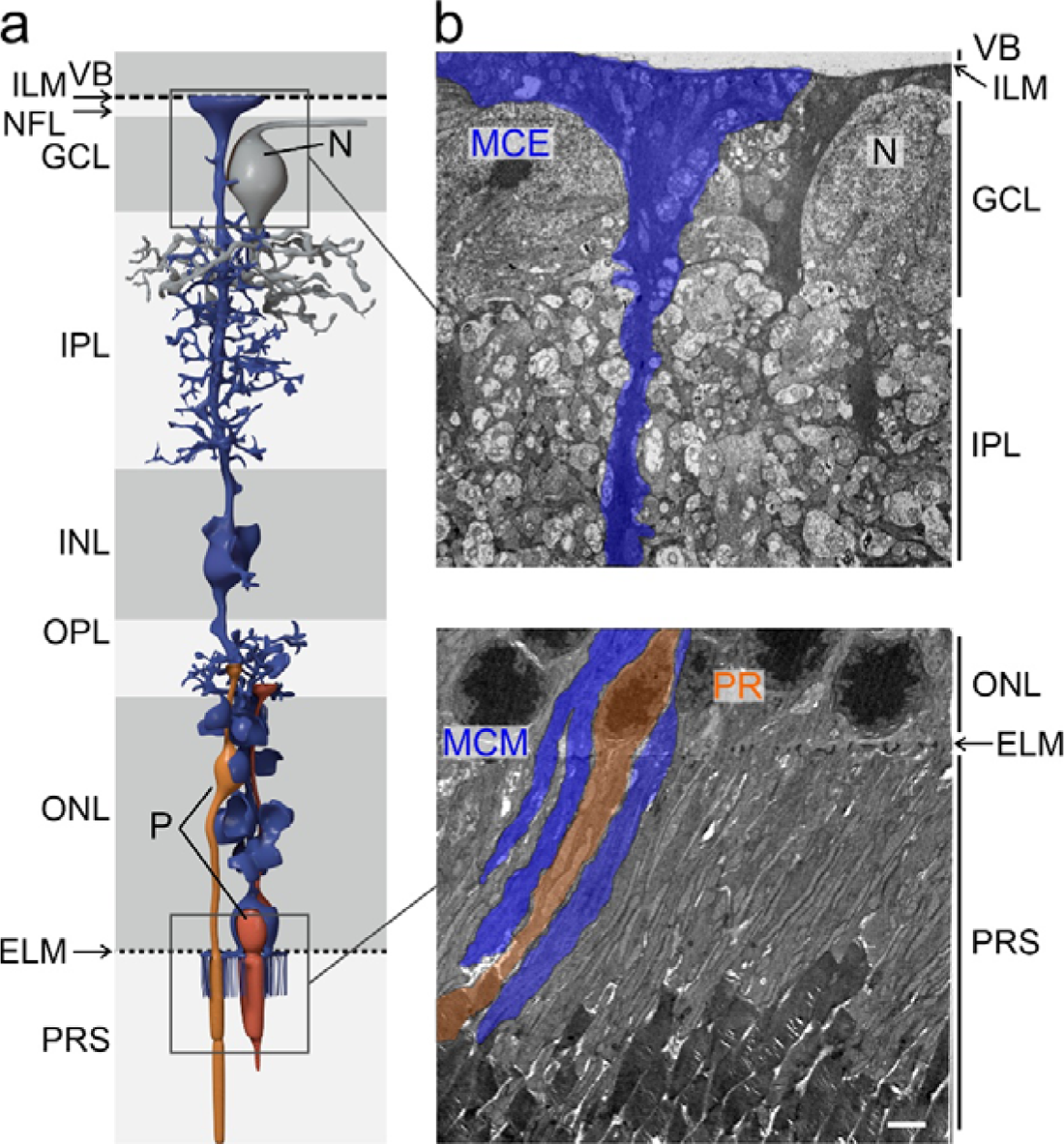
Location of Müller cells and ultrastructure of selected compartments in the inner and outer retina. a) Schematic representation of a Müller cell and its position with respect to the different layers of the mouse retina. VB, vitreous body; ILM, internal limiting membrane; NFL, nerve fiber layer; GCL, ganglion cell layer; N, soma of a neuron; IPL, inner plexiform layer; INL, inner nuclear layer; OPL, outer plexiform layer; ONL, outer nuclear layer; ELM, external limiting membrane; P, soma of a photoreceptor; PRS, photoreceptor segments. b) Representative transmission electron micrographs showing layers of the inner (top) and outer (bottom) retina and the location of Müller cell elements. In the inner retina (top), endfeet of Müller cells (MCE) face the vitreous body (VB), form the inner limiting membrane (ILM) and surround large somata of neurons (N) in the ganglion cell layer (GCL). Their processes traverse and interweave the inner plexiform layer (IPL; top). In the outer retina (bottom), Müller cell processes traverse the outer nuclear layer (ONL) containing nuclei (P) of photoreceptors (PR) and send apical microvilli (MCM) from the external limiting membrane (ELM) into the subretinal space where they interweave with photoreceptor segments (PRS). Scale bar: 2 µm.

**Figure 4.**
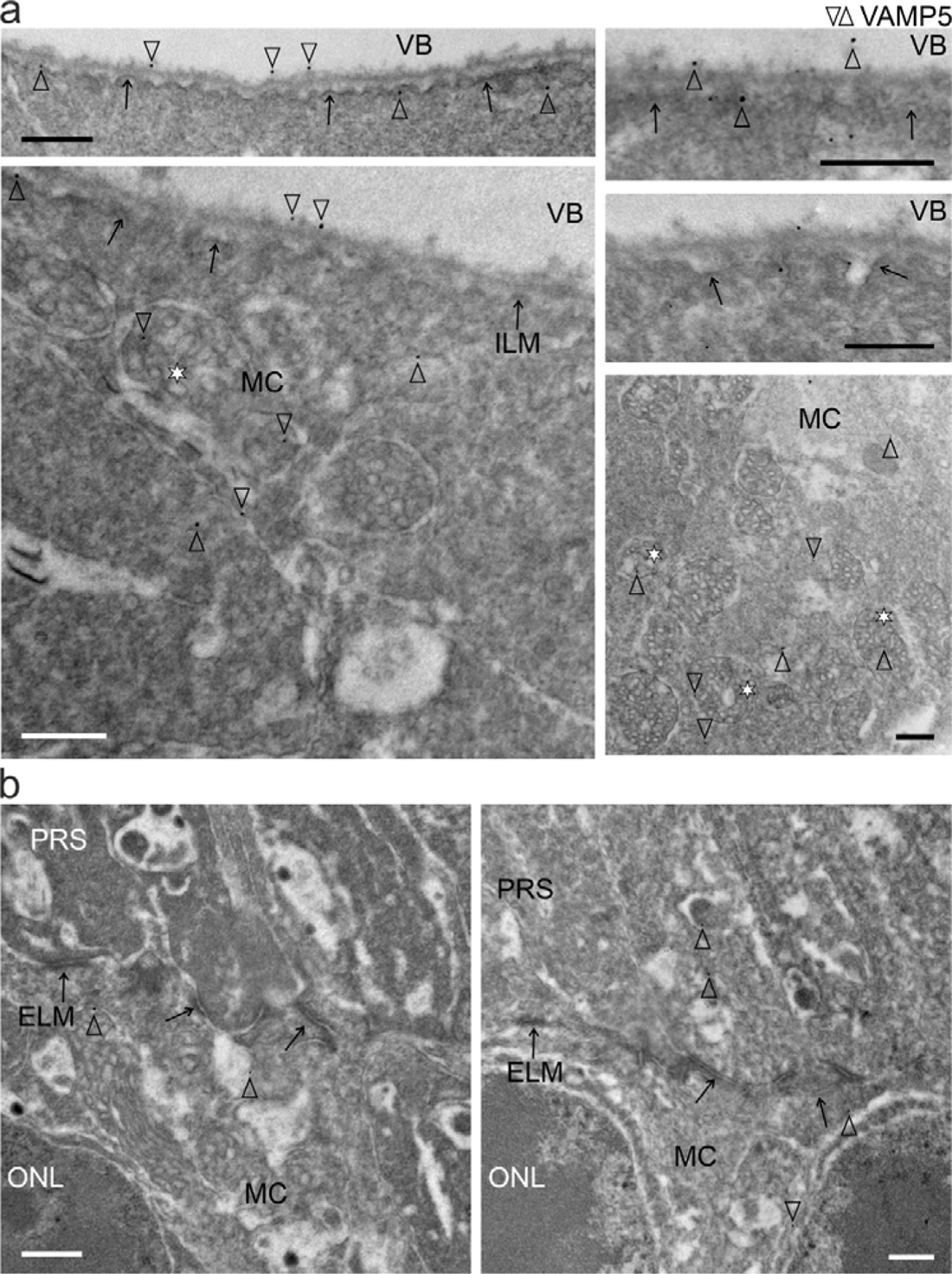
Subcellular and extracellular location of VAMP5 in the retina. (a) Transmission electron micrographs of the inner retina showing the presence of VAMP5 (empty arrowheads) in endfeet of Müller cells (MC), notably in multivesicular bodies (asterisks), at the intra- and extracellular side of the internal limiting membrane (ILM, black arrows) and in the vitreous body (VB). (b) Electron micrographs of the outer retina showing the presence of VAMP5 in processes of Müller cells (MC) in the outer nuclear layer (ONL) and on their apical microvilli extending beyond the external limiting membrane (ELM, black arrows) into the subretinal space containing photoreceptor segments (PRS). Retinal sections were subjected to immunogold labeling of VAMP5. Selected nanogold particles are indicated by empty arrowheads. Scale bars: 500 nm.

### Presence of EV markers in specialized domains of retinal Müller cells

Our ultrastructural findings raised the question whether VAMP5 is associated with EVs secreted by Müller cells. To explore this, we studied the distribution of CD81 (TSPAN28), CD9 (TSPAN29) and CD63 (TSPAN30) in Müller cells. These transmembrane proteins are ubiquitously expressed members of the tetraspanin family, located in distinct cellular compartments and contained in distinct types of EVs (Escola et al., 1998; Théry et al., 1999; Théry et al., 2018). Immunohistochemical and -cytochemical detection of the selected tetraspanins in retinal sections (Fig. 5a) and in acutely isolated immunoaffinity-purified Müller cells (Fig. 5b), respectively revealed a distinct distribution of each protein along GLUL-positive Müller cells. The cell surface component CD9 decorated their endfeet facing the vitreous body, structures in the outer plexiform layer and notably their apical microvilli extending into the subretinal space (Fig. 5a). CD81 was strongly expressed in the outer plexiform layer and on apical microvilli of Müller cells. CD63, which is also present in the endosomal-lysosomal system (Escola et al., 1998; Kobayashi et al., 2000; Pols and Klumperman, 2009), showed a distinct distribution. This protein was present in the endfeet of Müller cells and nearly absent from their apical microvilli (Fig. 5a). Inspection of acutely isolated Müller cells showed limited overlap of CD9 and CD81 and some colocalization of CD9 and CD63 (Fig. 5b). CD9 and CD63 also showed punctate staining not associated with GLUL (Fig. 5a) raising the question whether these proteins were expressed by other types of retinal cells. To address this question, we compared expression levels between retinal cell types using transcriptome (RNAseq) and proteome (mass spectrometry) analyses of acutely isolated and immunoaffinity-purified cells (Fig. 6). For each tetraspanin tested, transcript and protein levels were highest in Müller cells compared to other cells with the notable exception of CD81, which was also abundant in neurons (Fig. 6). Taken together, these results revealed a domain-specific presence of the selected tetraspanins with CD63 expressed on Müller cell endfeet facing the vitreous body, and CD9 and CD81 strongly present on apical microvilli of Müller cells extending into the subretinal space.

**Figure 5.**
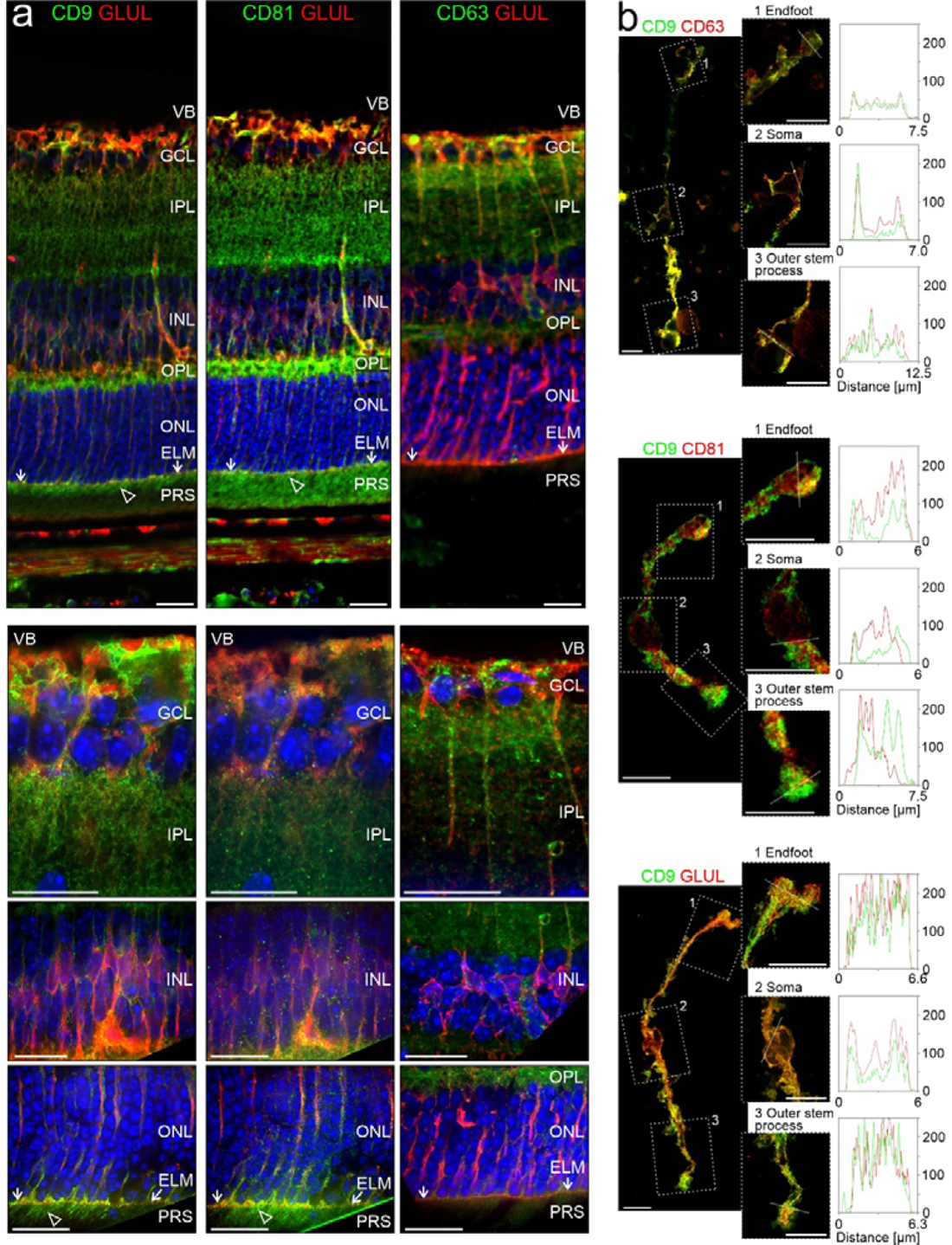
Distribution of EV markers in the retina. (a) False-color confocal micrographs of retinal sections subjected to double-immunohistochemical staining of GLUL-positive retinal Müller cells and of the tetraspanins CD9, CD81 or CD63. Empty arrowheads indicate presence of CD9 and CD81 on apical microvilli of Müller cells extending beyond the external limiting membrane (ELM, white allows) in the subretinal space containing photoreceptor segments (PRS). VB, vitreous body; GCL, ganglion cell layer; IPL, inner plexiform layer; INL, inner nuclear layer; OPL, outer plexiform layer; ONL, outer nuclear layer; RPE, retinal pigment epithelium. Scale bars: 20 µm. (b) False-color confocal (left) and STED (right) micrographs of acutely isolated, immunoaffinity-purified Müller cells showing entire cells and selected compartments, indicated by rectangles, at higher magnification. The terms “endfoot” and “outer stem process” indicate the Müller cell compartments that are apposed to the internal and external limiting membrane in the intact retina, respectively. Fixed cells were subjected to double-immunocytochemical staining of CD9, CD63 or CD81 and of GLUL. Scale bars: 10 µm. Plots show fluorescence intensities [in analog-digital units] of respective proteins across the selected compartments with the location of the line scans indicated by white dashed lines.

**Figure 6.**
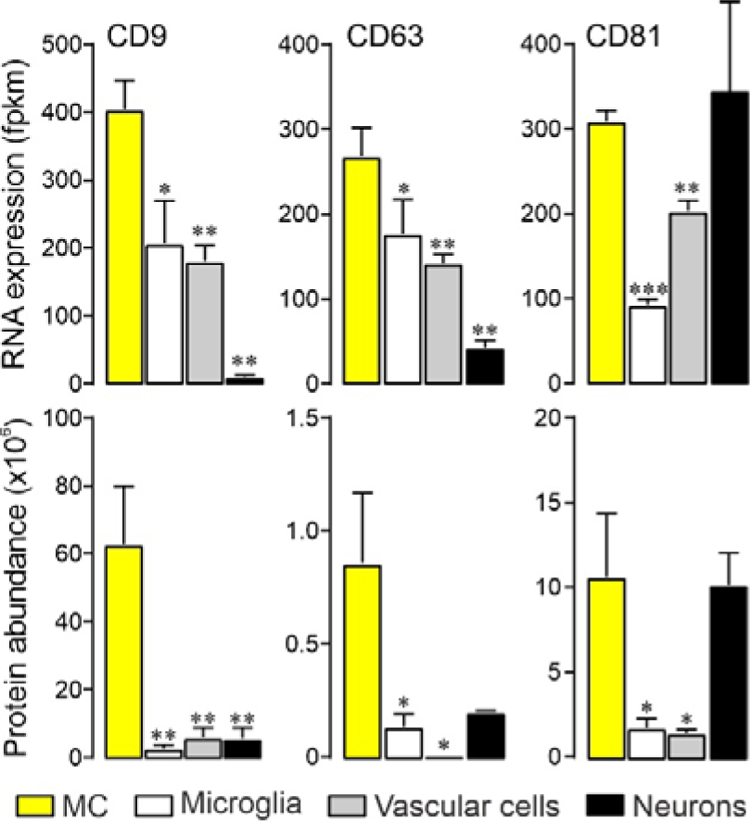
Cell-specific expression of tetraspanins in selected retinal cells. Transcript (top) and protein levels (bottom) of indicated tetraspanins in acutely isolated immunoaffinity-purified retinal cells (n = 3-5; whiskers: SEM; Mann-Whitney test) as determined by RNAseq and mass spectrometry, respectively. Fpkm, fragments per kilobase million. MC, Müller cells.

### Ultrastructural and molecular characterization of EVs secreted by Müller cells *in vitro*

Our observations *in vivo* suggested that Müller cells secrete EVs. Among the tetraspanins detected in the retina, we focused on CD63 and CD9 given their strong expression by glial cells, and their presence in Müller cell endfeet and microvilli representing highly specialized compartments of these cells. To characterize glia-derived EVs, we purified Müller cells from retinal cell suspensions by immunomagnetic separation, cultured them for maximally 60h under chemically defined, serum-free conditions and prepared secreted material contained in CM as outlined below. For comparison, we also analysed material in CM of affinity-purified retinal neurons. To determine cell viability during the indicated culture periods, we first performed TUNEL assays. This approach revealed absence of TUNEL-positive in GLUL-positive Müller cells indicating high viability of Müller cells. Among GLUL-negative cells representing in part neurons, a fraction of cells was TUNEL-positive indicating a lower, but stable viability of these cells during the culture period (Fig. 7a). Next, we characterized material secreted by retinal Müller cells and neurons by established methods (Théry et al., 2018). As first step, we enriched EV-like material contained in Müller cell- and neuron-derived CM by differential ultracentrifugation (Théry et al., 2006) and inspected its form and size by SEM. We observed round, vesicle-like structures (Fig. 7b) with diameters between 30 to 150 nm (Fig. 7c, top panel), which is within the established size range of EVs. Independent information about the size of these structures was obtained by NTA, which also informed about particle concentrations (Fig. 7c, bottom panel). To this end, we enriched material secreted by cultured Müller cells and neurons by immunomagnetic separation of CM using an antibody against CD9. These samples showed a narrower size range than the ultracentrifugation-enriched material inspected by SEM probably due to the immunoaffinity-based selection (Fig. 7c). Notably, NTA revealed a ∼1000-fold higher concentration of particles in affinity-purified samples compared to CM thus indicating efficient enrichment of EV-like material (Fig. 7c, insert). We next probed whether Müller cells secrete VAMP5. To this end, we purified CD63- and CD9-positive material from Müller cell- and neuron-derived CM and subjected the samples to immunoblotting. Flow-through lacking EVs and cell or retinal lysates served as negative and positive controls, respectively. This approach revealed the presence of VAMP5 – albeit at low levels – in CD9-positive material secreted by Müller cells, but not in material secreted by neurons and not in CD63-positive material from either cell population probably due to limited quantities (Fig. 7d). To test directly whether VAMP5 is present on EV-like structures, we subjected CD9- and CD63-positive material secreted by cultured Müller cells to double-immunogold staining followed by SEM or TEM. Electron microscopic inspection revealed the colocalization of VAMP5 with CD63 and with CD9 on EV-like structures (Fig. 7e,f).

**Figure 7.**
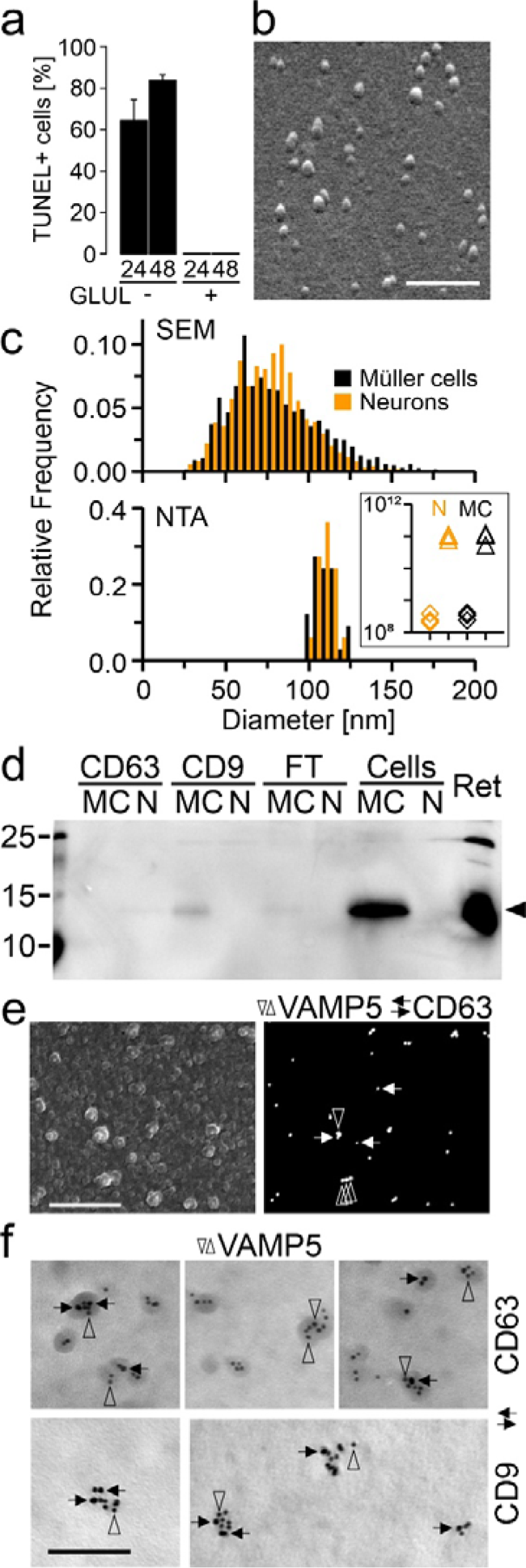
Characterization of EVs secreted by Müller cells *in vitro*. (a) Cell death among GLUL-positive Müller cells and GLUL-negative neurons after indicated periods (in hours) in serum-free culture estimated by TUNEL staining. Note the absence of TUNEL-positive cells in cultures of GLUL-positive cells. (b) Scanning electron micrograph of vesicle-like structures released by immunoaffinity-purified Müller cells cultured for 60h under serum-free conditions. Secreted material contained in CM was enriched by ultracentrifugation prior to processing for SEM. Scale bar: 600 nm. (c) Diameters of vesicle-like structures shown in (b) as determined by SEM (top; n = 3091 / 2361 particles from neurons / Müller cells; n = 3 culture preparations each; 10 images each preparation) and of CD9-positive EVs determined by NTA (bottom; n = 3 independent cultures). Orange and black bars indicate neuronal and glia-derived material, respectively. Insert, particle concentration per mL in CM (diamonds) and immunoaffinity-purified EV samples (triangles) from neurons (N, orange) and Müller cells (MC, black). (d) Immunoblots probing the presence of VAMP5 in CD63- or CD9-positive EVs secreted by cultured Müller cells (MC) and neurons (N), in the flow-through (FT), in cell lysates (Cells) and in the retina (Ret). (e) Scanning electron micrographs of secondary (left) and backscatter electrons (right) showing the surface and proteins, respectively of EVs immunoaffinity purified from CM of Müller cells and subjected to double-immunogold labeling of VAMP5 (vertical empty arrowheads; 40 nm) and CD63 (horizontal black arrows; 25 nm) using gold nanoparticles of different sizes. Scale bar: 750 nm. (f) Transmission electron micrographs showing colocalization of indicated proteins on affinity-purified EVs secreted by Müller cells *in vitro*. Ultrathin sections were subjected to double-immunogold labeling of VAMP5 (vertical empty arrowheads) and of CD63 or CD9 (horizontal black arrows) using gold nanoparticles of different sizes (VAMP5/CD9: 5/10 nm; VAMP5/CD63: 10/5 nm). Scale bar: 100 nm.

### Distinct proteomic profiles of CD9- and CD63-positive EVs secreted by neurons and Müller cells *in vitro*

We next determined the protein composition of immunoaffinity-purified material secreted by Müller cells and neurons in serum-free primary cultures using label-free mass spectrometry (Fig. 8). As controls, we analysed the protein content of non-binding flow-through of CM and of lysates of cultured and of acutely isolated cells, each from Müller cells and neurons (Fig. 8). The mass spectrometric data revealed an enrichment of established (CD9, CD81, ALIX, FLOT1, ANXA2) and of more specific EV components (COL1A2: Velázquez-Enríquez et al., 2021; LGALS3BP: Luga et al., 2012) in the CD9- and CD63-affinity purified samples compared to flow-through of media and to cell lysates (Fig. 8). On the other hand, proteins of organelles unrelated to EVs were only present in lysates, but absent from secreted material (Fig. 8). This observation excluded the presence of contaminating material and indicated the quality of our EV preparations. Glial and neuronal markers were enriched in the respective lysates confirming their cell-specific origin (Fig. 8). Among these proteins, the cytoplasmic components GLUL and MAP1B were also present in material immunoaffinity-purified from CM (Fig. 8). To explore the cell- and subtype-specific protein composition of presumed EVs, abundance values of proteins were subjected to hierarchical clustering (Fig. 9). The resulting sample clusters (columns) recapitulated the experimental groups indicating distinct compositions of neuron- and glia-secreted material and differences among CD9- and CD63-positive EV samples. The approach assigned proteins detected in EV samples (rows) to four principal groups (Fig. 9): Cluster 1 and cluster 2 contained either Müller cell- or and neuron-specific components, respectively. Two additional clusters contained proteins that were expressed in both neurons and Müller cells at high levels (Cluster 3) and at distinct levels (Cluster 4) across all experimental groups. Notably, all clusters contained EV components as shown by GO term analysis (Fig. 9). To further explore the diversity of glia- and neuron-derived EVs, we compared the protein contents of CD9- and CD63-positive material in CM from Müller cells and from neurons. As shown in Fig. 10a, between 59% to 64% of proteins detected in EV-like structures secreted by Müller cells or neurons were non-overlapping and thus specific to the cell of origin. The fraction was similarly large in CD9- and CD63-positive material. This finding indicated that neurons and Müller cells secrete EVs with partially distinct protein contents. A comparison of proteins in CD9- and CD63-positive EVs from neurons and Müller cells revealed 80% and 60% overlap, respectively (Fig. 10b) indicating that the two populations of EVs secreted by glial cells have a more distinct protein content than those secreted by neurons. A comparison of our proteomic data with entries in the Vesiclepedia database (Pathan et al., 2019) revealed that half of the proteins detected in neuron- and Müller cells-derived EVs are established markers (Fig. 10). Notably, we found CD81 and diazepam binding inhibitor (DBI) exclusively in CD9-positive EVs from Müller cells, whereas the fibrinogen beta chain (FGB) was only detected in their CD63-positive counterparts. The EV component galectin 3 binding protein (LGALS3BP) was present in neuronal and Müller cell-derived CD9-positive EVs (Fig. 10). Among the new proteins strongly enriched in Müller-derived EVs are centrosomal protein 290 (CEP290) and carbonic anhydrase (CA3). Taken together, our findings revealed that immunoaffinity-purified Müller cells secrete EVs *in vitro* that have a distinct protein composition from those secreted by neurons. Moreover, CD9- and CD63-positive EVs from Müller cells show limited protein overlap indicating distinct subsets.

**Figure 8.**
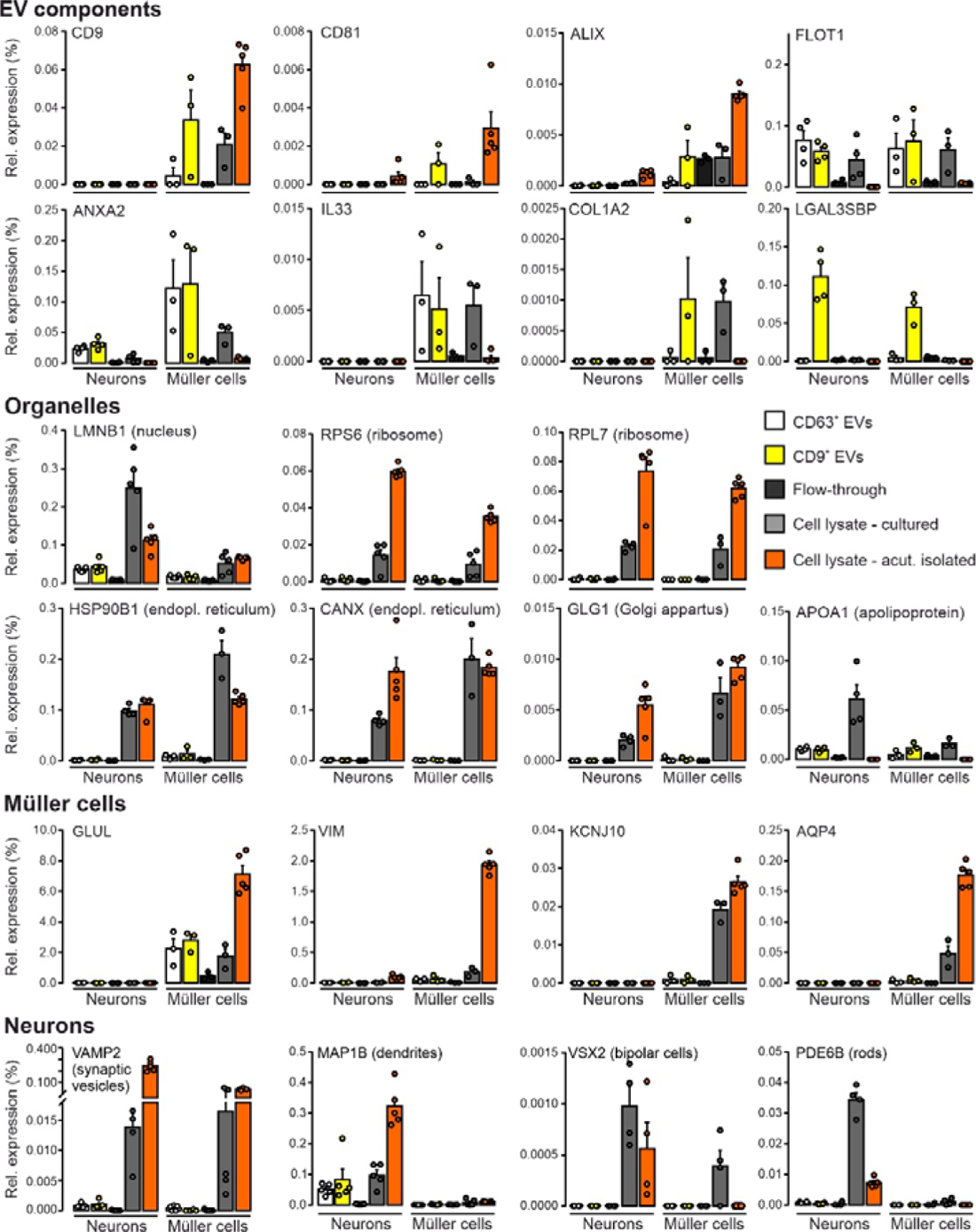
Presence of established markers in EVs secreted by Müller cells and neurons *in vitro*. Relative expression levels of selected proteins contained in EVs, organelles, Müller cells and neurons in CD9- and CD63-positive EVs purified from CM, in material of the flow-through, and in lysates of cultured and of acutely isolated neurons and Müller cells determined by mass spectrometry. Protein levels were normalized to total abundance of protein detected in respective samples. Symbols represent individual preparations. Bars and whiskers represent average values and SEM, respectively (n = 5 preparations).

**Figure 9.**
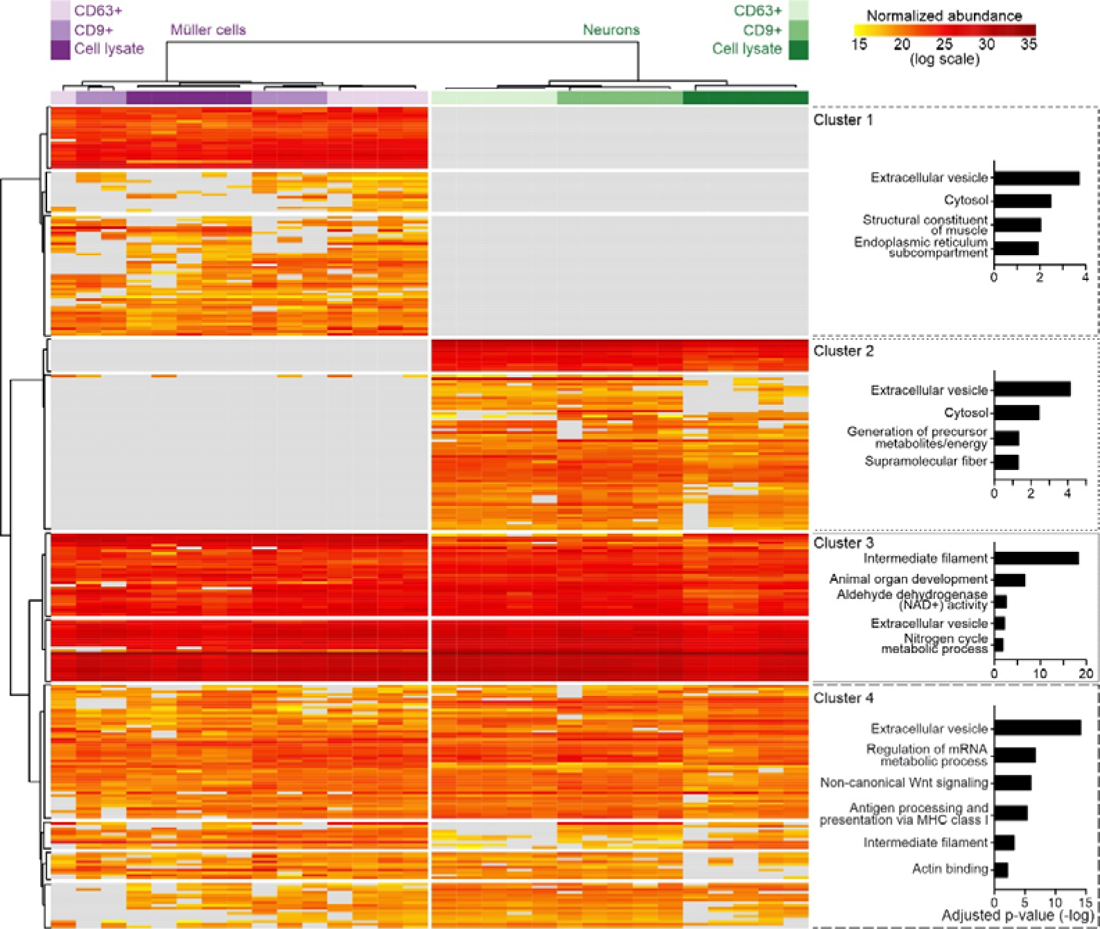
Protein profiles of CD9- and CD63-positive EVs from Müller cells and neurons. Heatmap showing the normalized abundances of proteins detected in EVs by mass spectrometry and dendrograms showing grouping of columns (experimental conditions) and of rows (proteins) generated by unsupervised hierarchical clustering. Plots on the right show GO terms in four largest protein clusters and respective adjusted p values.

**Figure 10.**
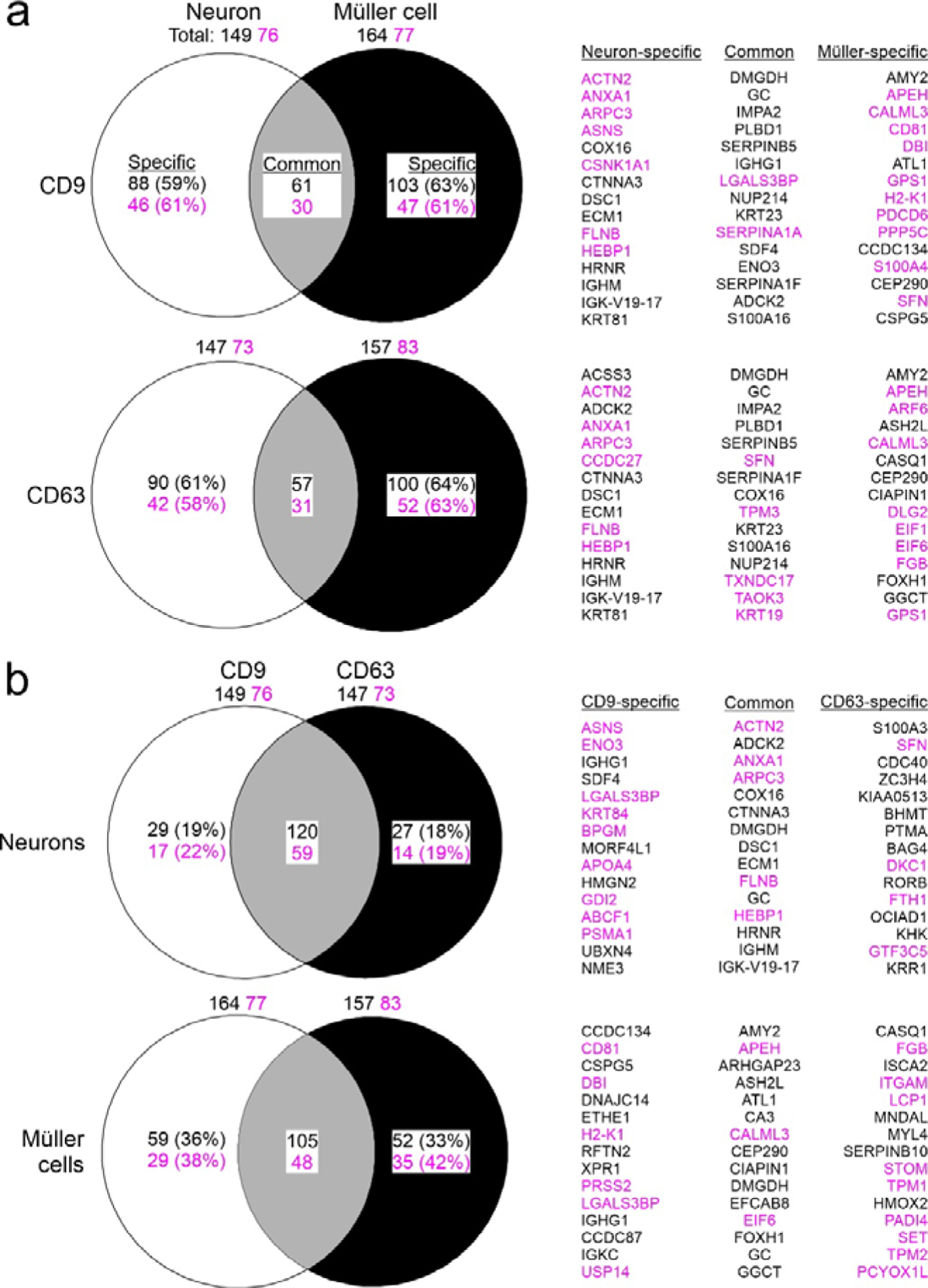
Diversity of neuron- and Müller cell-derived EVs based on protein content. Venn diagrams showing differences and overlaps in protein content between neurons and Müller cells for CD9-(top) and CD63-positive (bottom) EVs (a) and between CD9- and CD63-positive EVs from neurons (top) and from Müller cells (bottom) (b). Numbers indicate counts of all proteins that were detected in respective samples (total), that were present in both samples (common) and that were only present in either sample (specific). Indicated percentages were calculated compared to total number of detected proteins. Names of specific and common proteins showing highest enrichment in EVs compared to lysate (top 15) are indicated on the right. Numbers and names in magenta highlight established EV components listed in the Vesiclepedia database.

### Colocalization of VAMP5 with tetraspanins in the extracellular space of the inner and outer retina *in vivo*

Our finding that VAMP5 and selected tetraspanins are present on EV-like material secreted by Müller cells *in vitro* prompted us to examine their colocalization on extracellular structures in retinae *in vivo* using light and electron microscopy. Immunohistochemical staining of retinal sections revealed an overlapping distribution of VAMP5 with CD63 at Müller cell endfeet (Fig. 11a) and with CD9 in apical microvilli (Fig. 11b). The apposition of CD63 and VAMP5 in Müller cell endfeet was corroborated by super-resolution light microscopy (Fig. 11c,d). The post-ischemic increase of VAMP5 expression (Fig. 2) prompted us to test whether induction of transient ischemia affected the distribution of VAMP5 and CD9 at two distinct time points. Previous reports showed already an increase of retinal CD9 at 7 days post-injury (Vazquez-Chona et al., 2004; Iwagawa et al., 2020) similar as reported here for VAMP5 after induction of ischemia (Fig. 2). Ischemia induced time-dependent changes in the distribution of CD9 and VAMP5 in layers of the inner (Fig. 12a) and outer (Fig. 12b) retina. CD9 showed a more increased presence on plasma membranes, especially in apical microvilli of Müller cells, whereas VAMP5 showed a more wide-spread expression in post-ischemic retinae (Fig. 12).

**Figure 11.**
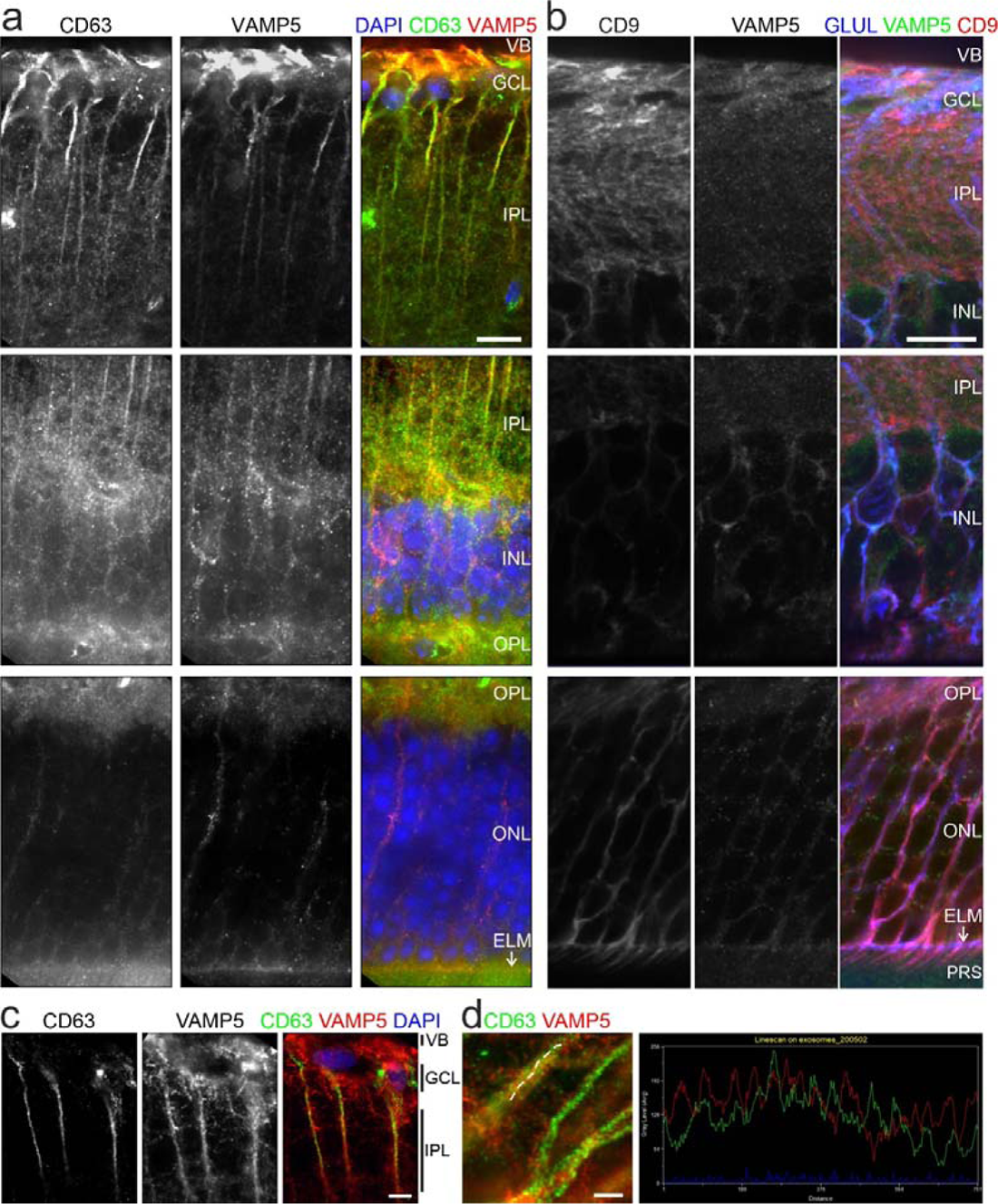
Colocalization of VAMP5 and EV markers in retinal cells *in vivo*. Greyscale and false-color confocal micrographs of retinal sections subjected to double-immunohistochemical staining of VAMP5 and of CD63 (a) or CD9 (b). VB, vitreous body; GCL, ganglion cell layer; IPL, inner plexiform layer; INL, inner nuclear layer; OPL, outer plexiform layer; ONL, outer nuclear layer; ELM, external limiting membrane (white arrows); PRS, photoreceptor segments. Scale bars: 10 µm. (c) Greyscale and false-color STED micrographs of retinal sections after double-immunohistochemical staining of CD63 and VAMP5. Scale bar: 5 µm. (d) STED micrographs at higher magnification showing Müller cell processes (left). Plot of fluorescence intensities along the line scan indicated in the micrograph (dashed white line) (right). Scale bar: 5 µm.

**Figure 12.**
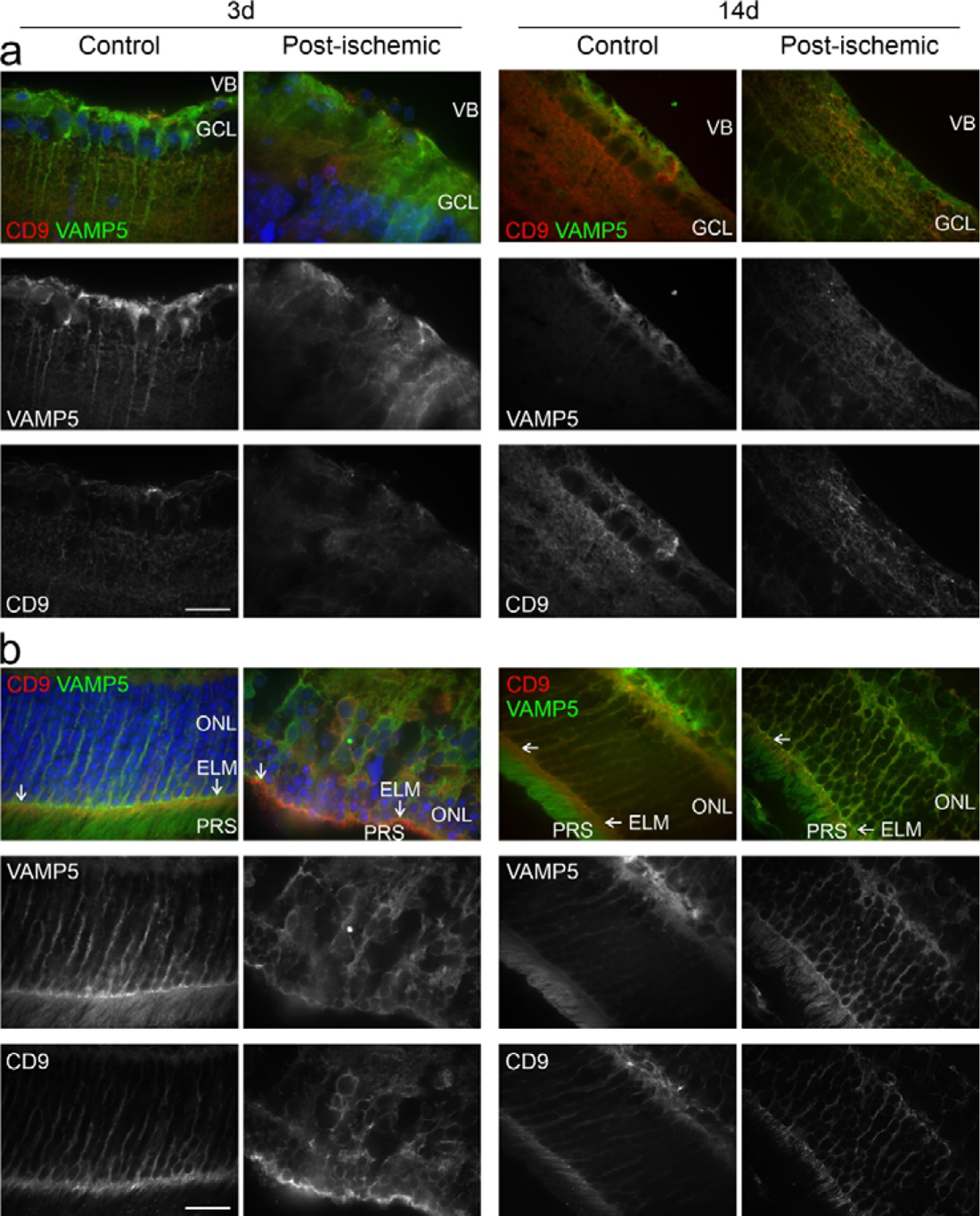
Post-ischemic changes in the retinal distribution of VAMP5 and CD9. False-color and corresponding greyscale confocal micrographs showing the inner (a) and outer (b) retinae from control eyes and from post-ischemic eyes at 3 (left) and 14 (right) days after transient ischemia. Retinal sections were subjected to double-immunohistochemical staining of VAMP5 (green) and of CD9 (red). VB, vitreous body; GCL, ganglion cell layer; ONL, outer nuclear layer; ELM, external limiting membrane (white arrows); PRS, photoreceptor segments. Scale bar: 40 µm.

Our light microscopic observations indicated overlap of VAMP5 and of tetraspanins in specialized domains of Müller cells situated in the inner and outer retina, but we could not distinguish intra- and extracellular structures. To address this crucial point, we performed double-immunogold labeling of respective proteins in ultrathin retinal sections using gold nanoparticles of distinct sizes (to facilitate orientation see also Fig. 3b). TEM inspection revealed close apposition of VAMP5 and CD63 in the extracellular space forming the vitreous body suggesting their presence on vesicle-like structures (Fig. 13a). Notably, we observed regularly vesicle-like structures decorated with VAMP5 and CD9 in the extracellular space intercalated between photoreceptor segments and Müller cell microvilli at variable distance from the external limiting membrane (Fig. 13b).

**Figure 13.**
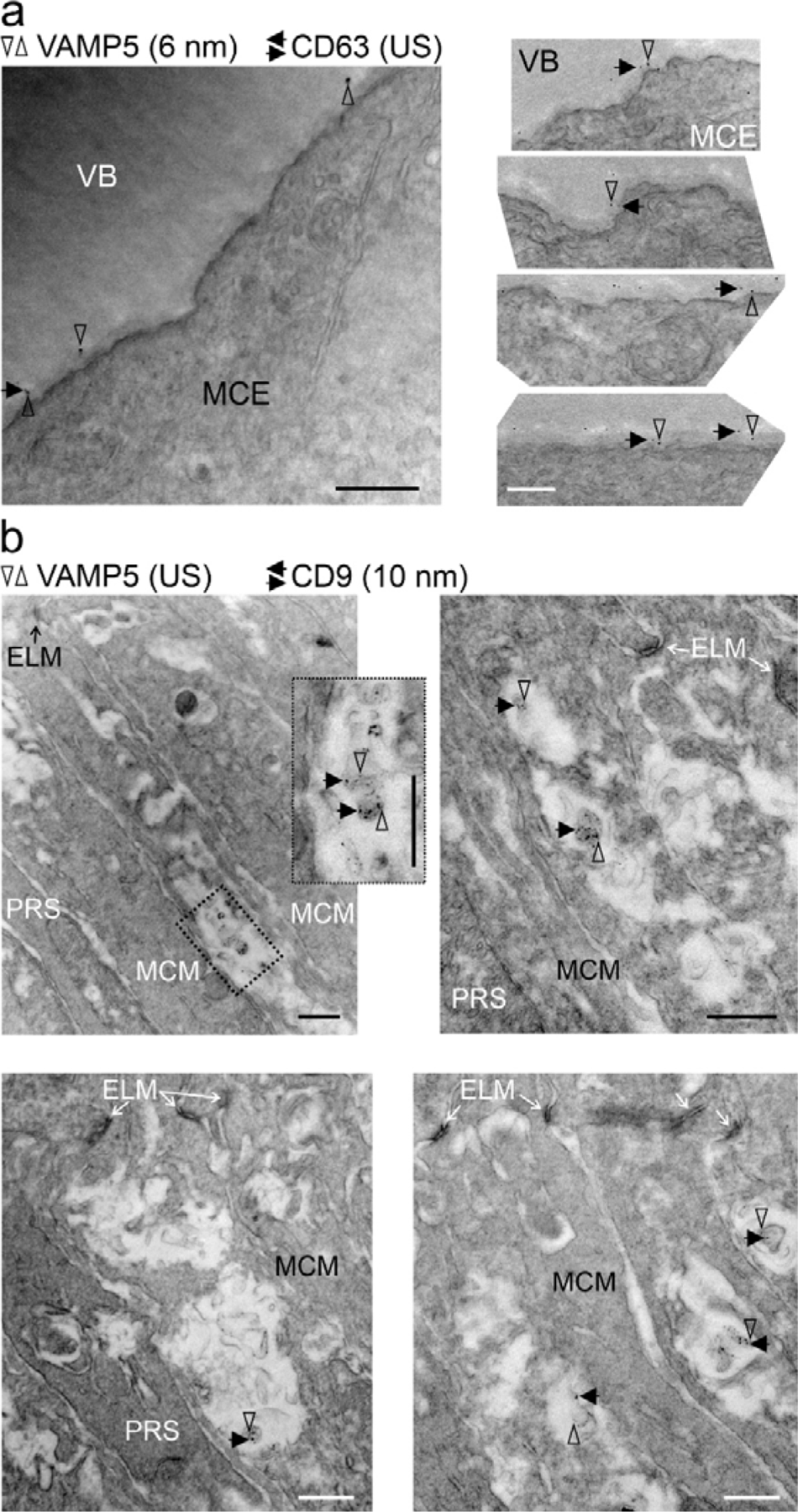
Colocalization of tetraspanins and VAMP5 on extracellular vesicular structures in the retina. Representative transmission electron micrographs of retinal sections revealing colocalization of CD63 (a) and of CD9 (b) with VAMP5 in the extracellular space (a) apposed to endfeet forming the vitreous body and (b) close to microvilli of Müller cells surrounding photoreceptor segments, respectively. Retinal ultrathin sections were subjected to double-immunogold labeling of the indicated proteins using gold nanoparticles of distinct sizes (as indicated; US, ultrasmall ∼ 0.8 nm). VB, vitreous body; MCE, Müller cell endfoot; ELM, external limiting membrane; MCM, Müller cell microvillus; PRS, photoreceptor segment. VAMP5: vertical empty arrowheads; CD63 and CD9: horizontal black arrows. Scale bars: 500 nm.

## Discussion

Our study provides first evidence that Müller cells release distinct EV-like structures from specialized compartments in the inner and outer retina of adult mice. Until now, EVs have mainly been studied after isolation from biological fluids or cell culture media. Here, we report their detection in an intact biological tissue *in vivo*, which has represented a key challenge in the field (van Niel et al., 2022; Verweij et al., 2021). Previously, EVs have been found in various ocular structures and fluids including acutely isolated retinae from mice (Mighty et al., 2020; Wooff et al., 2020), liquid biopsies of the human vitreous body (Zhao et al., 2018) and aqueous humor (Hsiao et al., 2021), the conjunctival mucin layer of rats (Tendler and Panshin, 2020), the retinal pigment epithelium of aged mice (Wang et al., 2009) and drusen of patients with age-related macular degeneration (Grillo et al., 2021). However, the cellular origin of these EVs remained unclear. Our findings point to Müller cells as a source of EVs in the retina and indicate that the release occurs at two prominent compartments of these cells. Müller cells are highly polarized showing distinct morphologic and functional specializations at their business ends: endfeet facing the vitreous body and apical microvilli surrounding photoreceptor segments (Derouiche et al., 2012; Reichenbach, 1989). Our results suggest that Müller cells secrete subsets of EVs from these compartments possibly by distinct mechanisms based on their tetraspanin content: The presence of CD63 in multivesicular bodies of endfeet and in the apposed extracellular space suggests that EVs in this compartment originate from intraluminal vesicles. The presence of CD9 on EVs in the subretinal space suggests that these structures originate from apical microvilli. Müller cell-derived EVs thus resemble exosome and ectosomes bearing CD63 and CD9, respectively. Their distinct subcellular origin and content have been exposed by a recent study on HeLa cell-derived material (Mathieu et al., 2021). Our hypothesis that Müller cells release EVs from their apical microvilli in the subretinal space is supported by the notion that many cell types shed EV-like structures from their specialized plasma membrane protrusions such as cilia, microvilli and filopodia (Marzesco et al., 2005; Hara et al., 2010; Wood et al., 2013; Wang et al., 2014a; Salinas et al., 2017; for review see Rilla, 2021). The presence of CD81 (Clarke and Geisert, 1998) and CD9 (Iwagawa et al., 2020) in Müller cell compartments of the inner and outer retina is supported by earlier reports. So far, the retinal distribution of CD63 has remained unknown. We report its enrichment in Müller cell endfeet. With respect to other retinal cells, this ubiquitously expressed tetraspanin has been studied in cultured human retinal pigment epithelial cells as part of their EVs (Sreekumar et al., 2010) and in rodent retinal ganglion cells as component of the endosomal-lysosomal system (Demais et al., 2016).

Until now, EVs secreted by primary Müller cells *in vitro* have not been characterized except for a recent report showing the presence of micro RNAs in these structures (Akamine et al., 2021). In fact, Müller cells have been mainly studied as target of EVs (Didiano et al., 2020; Eastlake et al., 2021; Kamalden et al., 2017; Ke et al., 2021; Peng et al., 2018; Wassmer et al., 2017; Zhang et al., 2020). Probably the best studied source of retinal EVs are cultured pigment epithelial cells (Ahn et al., 2021; Flores-Bellver et al., 2021; Kang et al., 2014; Klingeborn et al., 2017; Mukai et al., 2021; Otsuki et al., 2021; Sreekumar et al., 2010; Toyofuku et al., 2012). Few studies characterized EVs secreted by other types of retinal cells *in vitro* including precursor cells (Zhou et al., 2018), astrocytes (Hajrasouliha et al., 2013), retinal ganglion cells (Wang et al., 2021) and photoreceptors (Kalargyrou et al., 2021). Microvesicle release from photoreceptors was reported *in vivo* (Ropelewski and Imanishi, 2020). Here, we provide the first direct comparison of neuronal and glial secretomes and of the respective cell lysates. Our analysis reveals that Müller cells and retinal neurons secrete EVs with distinct protein compositions showing less than 40% overlap. We uncovered several Müller cell-specific EV components. DBI is a putative endogenous ligand of translocator protein 18 kDa (TSPO) thought to mediate interactions of Müller cells and microglia in the retina (Wang et al., 2014b). Carbonic anhydrase was detected on apical microvilli of Müller cells in the mouse retina (Nagelhus et al., 2005; Ochrietor et al., 2005) supporting the release of EVs from these structures. CEP290/NPHP6, a ciliary component, has not been associated with EVs, but its deficiency causes retinal diseases (Chen et al., 2021). EVs secreted by cultured neurons (Chivet et al., 2014; Faure et al., 2006; Lachenal et al., 2011; Morel et al., 2013; Yuyama et al., 2012) and glial cells (Frühbeis et al., 2013; Gabrielli et al., 2015; Glebov et al., 2015; Guitart et al., 2016; Hooper et al., 2012; Potolicchio et al., 2005) from other parts of the CNS have been characterized, but their proteomic profiles have not been compared. Notably, the limited overlap of proteins found in CD9- and CD63-positive EVs from Müller cells supports the notion that a given cell type produces distinct subtypes of EVs (Jeppesen et al., 2019; Kowal et al., 2016) and corroborates our idea that Müller cells secrete distinct types of EVs from distinct compartments.

Our analysis of VAMPs revealed Müller cell-specific expression of VAMP5. Up to now, the distribution of this protein in the retina has remained unknown. Previous reports detected VAMP5 in skeletal muscle and other organs, but not in the brain (Takahashi et al., 2013; Zeng et al., 1998). A more recent study identified *Vamp5* as miRNA-regulated target gene in the retina supporting our finding that the protein is present in this part of the CNS (Olivares et al., 2017). Our finding that VAMP5 is associated with EVs is supported by several lines of evidence. Electron microscopic inspection of muscle cell lines revealed its localization at the plasma membrane, in vesicular structures, and in multivesicular bodies (Tajika et al., 2014; Zeng et al., 1998) in line with our immunogold data. Moreover, VAMP5 was detected in EVs of the murine lung (Choudhary et al., 2021), from cultured myotubes (Forterre et al., 2014), mesenchymal stem cells (Salomon et al., 2013) and tumoral Jurkat T cells (Bosque et al., 2016). A hint towards the function of VAMP5 in EVs comes from a recent observation that components of the SNARE complex can be transferred intercellularly via EVs enabling vesicular release in target cells (Vilcaes et al., 2021). VAMP5 has also been shown to be part of the machinery that mediates EV-dependent cytokine release from mesenchymal stem cells *in vitro* (Kou et al., 2018). VAMP5 did not mediate fusion of vesicular and plasma membrane containing specific tSNAREs in an artificial exocytosis assay (Hasan et al., 2010), but deletion of VAMP5 from mice causes developmental defects in the respiratory and urinary systems and perinatal death (Ikezawa et al., 2018). The observed upregulation of VAMP5 and the changes in CD9 distribution under ischemic conditions suggest that VAMP5 is part of the gliotic reaction to injury and disease and that ischemia affects EV secretion in the retina. Ischemia-induced changes in brain cells releasing EVs have been reported previously (Brenna et al., 2020). On the other hand, EVs are explored as therapeutic approaches to mitigate post-ischemic pathologic changes (Doeppner et al., 2015).

In summary, our data support the idea of EV-based communication between cells in the CNS and suggest Müller cells as a key actor in the retina.

## Acknowledgements

*Author contributions*: VD: performed electron microscopy-related work, acquired and analysed electron micrographs. AP: performed immunofluorescence microscopy-related work, qPCR experiments and analyses, immunoblots for VAMP expression in the healthy and ischemic retina, established protocols for primary Müller cell cultures and performed EV purification. KW: performed EV purification from primary Müller cell cultures and immunoblot analyses, analyzed EV marker localization via STED microscopy. AMP: generated purified EVs for mass spectrometric analysis. LK: performed EV isolation and bioinformatic analysis of proteomic data. AB: performed immunohistochemical staining with selected antibodies to prepare immunogold staining. RD: performed NTA. BP: set up the EV isolation protocol, analysed results, edited the manuscript. BG: analysed NTA results and edited the manuscript. SMH: performed mass spectrometric analysis of retinal cell types and purified EVs, analysed protein expression levels and edited the manuscript. AG: designed experiments, analysed results, prepared figures and edited the manuscript. FWP: designed experiments, analysed results, prepared figures and wrote the manuscript. The authors thank Jens Grosche for the schematic drawing of retinal cells, and Alexandra Hauser, Dirkje Felder and Gabrielle Jäger for excellent technical assistance.

## Funding statement

Part of this work was supported by the Centre National de la Recherche Scientifique under grant UPR3212 (FWP), the Université de Strasbourg under grant UPR3212 (FWP), the Agence National de la Recherche Scientifique under grant GLIAVAMP (FWP), the Deutsche Forschungsgemeinschaft under grants GR 4403/1-1; GR 4403/5-1; GR 4403/7-1 (AG) and HA 6014/5-1 (SMH) and the ProRetina Foundation Germany under grants Pro-Re/Seed/Grosche.1-2014 (AG) and Pro-Re/Seed/Kaplan-Grosche.1-2019 (LK, AG).

## Declaration of Interest Statement

The authors report no conflict of interest.

## Data Availability Statement

Data are available on request from the authors.

## Notes

### Competing Interest Statement

The authors have declared no competing interest.

### Summary of Updates

Ms completely revised and corrected according to referees' suggestions. J Extracell Ves, in press.

