## Supplemental tables for "Release of VAMP5-positive extracellular vesicles by retinal Müller glia *in vivo*"

**Demais et al.**

**Supplementary table 1.** Primary and secondary antibodies.

| **Primary antibodies** | | | | |
| --- | --- | --- | --- | --- |
| **Target** | **Host** | **Company** | **Reference** | **Dilution** |
| GLUL | mouse | Merck (Darmstadt, Germany) | MAB302 | 1:1000 (IF) |
| CRALBP | rabbit | Santa Cruz (Dallas, USA) | sc-28193 | 1 : 200 (IF) |
| VAMP1 | rabbit | Synaptic Systems (Göttingen, Germany) | 104 002 | 1 : 100 (IF)  1 : 5000 (WB) |
| VAMP2 | mouse | Synaptic Systems | 104 211 | 1 : 100 (IF)  1 : 5000 (WB) |
| VAMP3 | rabbit | Synaptic Systems | 104 103 | 1 : 100 (IF)  1 : 5000 (WB) |
| VAMP4 | rabbit | Thermofisher Scientific (Schwerte, Germany) | PA1-768 | 1 : 100 (IF)  1 : 5000 (WB) |
| VAMP5 | rabbit | Synaptic Systems | 176 003 | 1 : 100 (IF)  1 : 5000 (WB)  1: 100 (EM) |
| VAMP7 | mouse | Synaptic Systems | 232 011 | 1 : 500 (IF)  1 : 5000 (WB) |
| VAMP8 | rabbit | Synaptic Systems | 104 302 | 1 : 100 (IF)  1 : 5000 (WB) |
| CD9 | rat | BD Biosciences (Heidelberg, Germany) | 553758 | 1 : 100 (IF)  1 : 500 (WB)  1 : 50 (EM) |
| CD63 | mouse | Abcam (Cambridge, UK) | ab108950 | 1 : 100 (IF)  1 : 1000 (WB)  1 : 50 (EM) |
| CD81 | rabbit | Cell Signaling Technology (Frankfurt, Germany) | 10037 | 1 : 100 (IF) |

| **Secondary antibodies** | | | | |
| --- | --- | --- | --- | --- |
| **Target** | **Host** | **Company** | **Reference** | **Dilution** |
| Rabbit IgG‐HRP | goat | Merck Millipore (Darmstadt, Germany) | 401315 | 1:10000 |
| Mouse IgG Ig-HRP | goat | Merck Millipore (Darmstadt, Germany) | 401215 | 1:10000 |
| Rabbit IgG | donkey | Thermofisher Scientific | A-21206 | 1:500 |
| Rat IgG | goat | Abberior (Göttingen, Germany) | 2-0132-005-1 | 1 : 200 |
| Mouse IgG | goat | Abberior | 2-0002-005-1 | 1 : 200 |
| Rat IgG- 10 nm gold particles | goat | Electron Microscopy Sciences/Euromedex, Souffelweyersheim, France | 25189 | 1:100 |
| Goat IgG – 15 nm gold particles | Donkey | Electron Microscopy Sciences | 25807 | 1:100 |
| Goat IgG – ultrasmall gold particles | Donkey | Electron Microscopy Sciences | 25801 | 1:100 |
| Rabbit IgG - 6 nm gold particles | Goat | Electron Microscopy Sciences | 25363 | 1:100 |
| Rabbit IgG - ultrasmall gold particles | Goat | Electron Microscopy Sciences | 25412 | 1:100 |

**Supplementary table 2.** Primer and TaqMan probe combinations for the detection of VAMPs by qRT-PCR.

| **Gene ID** | **Primer Sequences: *forward*** | **Primer Sequences: *reverse*** | **Roche TaqMan® Probe** |
| --- | --- | --- | --- |
| Vamp1 | 5‘ acg gac ctc cac ttc ctc tt 3‘ | 5‘ cag ctc ctt ctg tcc ctt ca 3‘ | # 78 |
| Vamp2 | 5‘ cca agc tca agc gca aat 3‘ | 5‘ggg att taa gtg ctg aag taa acg 3‘ | # 31 |
| Vamp3 | 5‘ttg aaa caa gtg ctg cca ag 3‘ | 5‘ atc cct atc gcc cac atc t 3‘ | # 31 |
| Vamp4 | 5‘tgc aag aga ata tta caa agg taa ttg 3‘ | 5‘ gaa agc ggt ggc att atc c 3‘ | # 91 |
| Vamp5 | 5‘ gag ttg gag cag cgt tca g 3‘ | 5‘ tgg gct aaa gtc ttg gtt gtc 3‘ | # 17 |
| Vamp7 | 5‘ gac ctt cgc ccc tca gtc 3‘ | 5‘ aga atg gcc atg act tca atc t 3‘ | # 78 |
| Vamp8 | 5‘ cct ccg aaa caa gac aga gg 3‘ | 5‘ gga caa tca cac aga tga tga ca 3‘ | # 31 |
